# Whole Blood Deep Immunophenotyping and Platelet-Leukocyte Aggregate Quantification by Spectral Flow Cytometry

**DOI:** 10.64898/2026.09.30.755388

**Authors:** Haiyuan Ding, Krista R. Gerbino, Katy Gilham, Yawei Tong, Alex Hu, Elaine Li Huang, Ella Zabriskie, Tahsin Ozpolat, Li Guo, Scott Brakenridge

**Author notes:** Co-corresponding authors: Li Guo, MD PhD, Department of Surgery, Harborview Medical Center, University of Washington School of Medicine, 300 9th Ave, Harborview Research & Training, Floor 6, Room 627, Seattle, WA, 98104; Scott Brakenridge, MD, Department of Surgery, Harborview Medical Center, University of Washington School of Medicine, 300 9th Ave, Harborview Research & Training, Floor 6, Room 621, Seattle, WA, 98104. **Data Sharing Statement:** For original data, please contact Dr. Guo or Dr. Brakenridge.

## Abstract

Platelet-leukocyte aggregates (PLAs) are a proposed interface between coagulation and immune regulation, but their distribution across leukocyte subsets beyond monocytes and neutrophils, and their behavior in acute disorders of immune regulation, such as sepsis and severe trauma, remain largely undefined. The study of thromboimmune mechanisms in these acute conditions has relied heavily on murine sepsis models, whose species differences in platelet and leukocyte biology limit translation to human disease. Direct measurement in human whole blood is needed yet reported PLA frequencies vary widely because quantification is highly sensitive to sample handling. Here we present a robust, minimally manipulative 27-color spectral flow cytometry protocol that allows simultaneous deep immunophenotyping and classification of PLAs across >40 leukocyte subpopulations. In healthy donors, PLA frequency ranged from 1.7% on memory CD4 T cells to over 50% on monocyte subsets, with monocytes contributing 25% of all PLAs. We also found novel platelet-binding leukocyte populations, including Th9 and Th22 CD4+ T cells. Optimization identified anticoagulant choice, centrifugation conditions, dilution, and sample processing time as key determinants of reproducible measurement; PLA quantification was validated by imaging flow cytometry. In a small validation cohort of sepsis and trauma patients, the assay recapitulated established immunophenotypic features of critical illness, including neutrophilia and myeloid-derived suppressor cells (MDSC) expansion, and quantified PLAs within each subset. This protocol provides a reference framework for studying thromboinflammation and other dysregulated immune response diseases directly in human blood, and a foundation for further studies, defining the mechanisms and clinical relevance of platelet-leukocyte interactions in disease.

**Key points:**

- Novel 27-color protocol enables simultaneous deep immunophenotyping and PLA quantification, validated by imaging flow cytometry.
- PLA measurements are highly sensitive to technical variables; Optimization of protocol conditions improves assay reproducibility.

## Introduction

Platelets and leukocytes have mainly been studied separately: platelets primarily in hemostasis and thrombosis, whereas leukocytes have largely been studied for inflammation and host immune responses.^1,2^ However, thrombosis, inflammation, and immune responses have been found closely interconnected.^3^ ‘Thromboinflammation’ was proposed to describe the integrated vascular responses that occur during microbial infection, vascular injury, thrombosis, and inflammation.^4,5^ Consistent with this concept, interactions between platelets and leukocytes, including neutrophils, monocytes, and lymphocytes, are increasingly recognized in both physiological and pathological settings.^3,5–9^

Ex vivo measurements of platelet–leukocyte aggregates (PLAs) in the peripheral blood could provide a snapshot of cell-cell interactions in vivo during thromboinflammation.^10–13^ Platelet-monocyte (PMAs), platelet-neutrophil (PNAs), and platelet-lymphocyte aggregates have been found increased in several diseases, including sepsis, COVID-19, AIDS, and acute coronary syndrome.^6,7,14–20^ Multiple adhesion receptors and receptor–ligand pairs are known to contribute to PLA formation, including P-selectin/PSGL-1 and GPIbα/αMβ2 integrin (CD11b/CD18; complement receptor 3 [CR3]; Mac-1).^8,21^ Although platelet-monocyte and platelet-neutrophil aggregates have been quantified in sepsis and after severe injury, the distribution of PLAs across the broader leukocyte subpopulations in these conditions remains undefined. Thus, the cellular composition, phenotypic signatures, and functional significance of PLAs in health and disease remain largely elusive. Across studies, the PLA baseline range in healthy populations, varies widely due to technical variations including anticoagulant choice, sample processing procedure, and processing time.^22–31^

Additionally, the study of thromboimmune mechanisms in these acute conditions has relied heavily on murine models. However, between human and mouse, there are significant differences in leukocyte constitution, platelet number and function, as well as their responses against microbial infections and traumatic injuries.^32^ For example, murine sepsis and trauma models diverge from human disease in transcriptomic, temporal, and innate and adaptive immune response patterns.^33–35^ Direct measurement in human whole blood is therefore required to define PLA biology that is relevant to patients.^34,36,37^

Recent advances in full-spectrum flow cytometry provide an opportunity for comprehensive profiling of PLAs.^38–40^ In this study, we designed a 27-color spectral flow cytometry panel and accompanying protocol capable of deep immunophenotyping of PLAs, across both major and minor myeloid and lymphoid compartments. By including >40 different leukocyte subpopulations together with several function markers, we found marked heterogeneity of PLAs across leukocyte populations, and among individuals. We further assessed technical variables, including anticoagulant choice and sample handling conditions, while validating inter-operator differences, day effects, and intra-operator effects. Our approach could provide a reference framework for platelet–leukocyte aggregate analysis in many diseases and may support the identification of disease-associated cellular signatures with diagnostic or prognostic potential.

## Methods

### Human subjects and Specimen collection

This study comprised a spectral flow cytometry discovery cohort with three population groups: healthy controls (HC), sepsis patients (SP), and trauma patients (TR). Healthy controls enrolled had not taken any known platelet-altering medication within the past 14 days. The sepsis samples were collected between days 2 and 3 of sepsis diagnosis according to Sepsis-3 Criteria (aged between 61 and 68). Trauma samples were collected from patients with an ISS score of 25 or greater and between days 2 and 5 post incident (aged between 22 and 75). Informed consent was obtained from the patient, or proxy if incapacitated. This study was approved by the University of Washington Institutional Review Board (IRB), with written informed consent obtained.

Venous blood was drawn sequentially into sodium citrate tubes (CIT) (BD #366560), sodium heparin tubes (HEP) (BD #367878), K_2_EDTA tubes (EDTA) (BD #367841), and acid citrate dextrose tubes (ACD) (BD #364606), in the respective order. Unless otherwise specified, all samples were processed within 0.5 hours of phlebotomy. Blood collected in EDTA tubes was used for complete blood count with differential (CBC/diff) analysis via Beckman Coulter’s DxH 500 Hematology Analyzer.

### Spectral flow cytometry

Fresh whole blood (100μL) was aliquoted into 5 mL flow tubes and incubated with TruStain FcX (Biolegend) and Live/Dead Blue at room temperature for 10 min, then stained with antibody mixture prepared with Brilliant Staining Buffer (BD Horizon) and Cellblox Blocking Buffer (Fisher) at room temperature for 30 min in dark. eBioscience™ 1-step Fix/Lyse Solution (Invitrogen) was then used to lyse red blood cells and fix stained samples. After centrifugation, cell pellets were resuspended in eBioscience™ Flow Cytometry Staining Buffer (Invitrogen) and stored at 4°C protected from light. Flow cytometry data were collected on a 5-laser Cytek Aurora spectral cytometer (Cytek Biosciences) and analyzed using FlowJo (version 10.10.0). Unless specified, data were collected after overnight storage at medium sample flow rate (∼30 μL/min). To evaluate effects of storage time (time to flow), duplicated samples were stored for 3 days post-fixation before data acquisition.

#### Centrifugation effects

Centrifugation conditions of 50 × g for 10 min, 100 × g for 10 min, 200 × g for 10 min, and 400 × g for 5 min were compared. Blood samples from the same healthy donor were centrifuged under each condition. Cell pellets were resuspended in Staining Buffer. CBC/diff analyses were performed on the resuspended pellet fractions using a DxH 500 hematology analyzer to assess the efficiency of free platelet removal from leukocyte pellets. To evaluate the impact of centrifugation on downstream flow cytometric measurements, in indicated experiments, samples were stained and lysed/fixed, then centrifuged at either 50 × g or 200 × g conditions followed by overnight storage and acquisition on the Aurora cytometer.

#### Dilution effects

The effects of sample dilution at multiple stages of processing were evaluated, including (i) dilution of whole blood 2-, 4-, or 8-fold prior to staining; (ii) dilution during fixation/lysis, either 1,000 μL or 2,000 μL of Fix/Lyse reagent was added; and (iii) dilution of fixed samples 2-, 4-, or 8-fold immediately before acquisition on the Cytek Aurora.

#### Time-to-stain effects

The impact of delayed staining was evaluated by staining samples either immediately after collection (within 0.5 hours of blood draw), 2 h post-collection, or 4 h post-collection.

### Data Analysis

Flow cytometry data were analyzed using FlowJo software (v10.10.0) according to the gating strategy shown in Figures 1A and 1B. Cell counts and the percentage of CD41+ for each leukocyte population were exported for downstream analysis.

**Figure 1:**
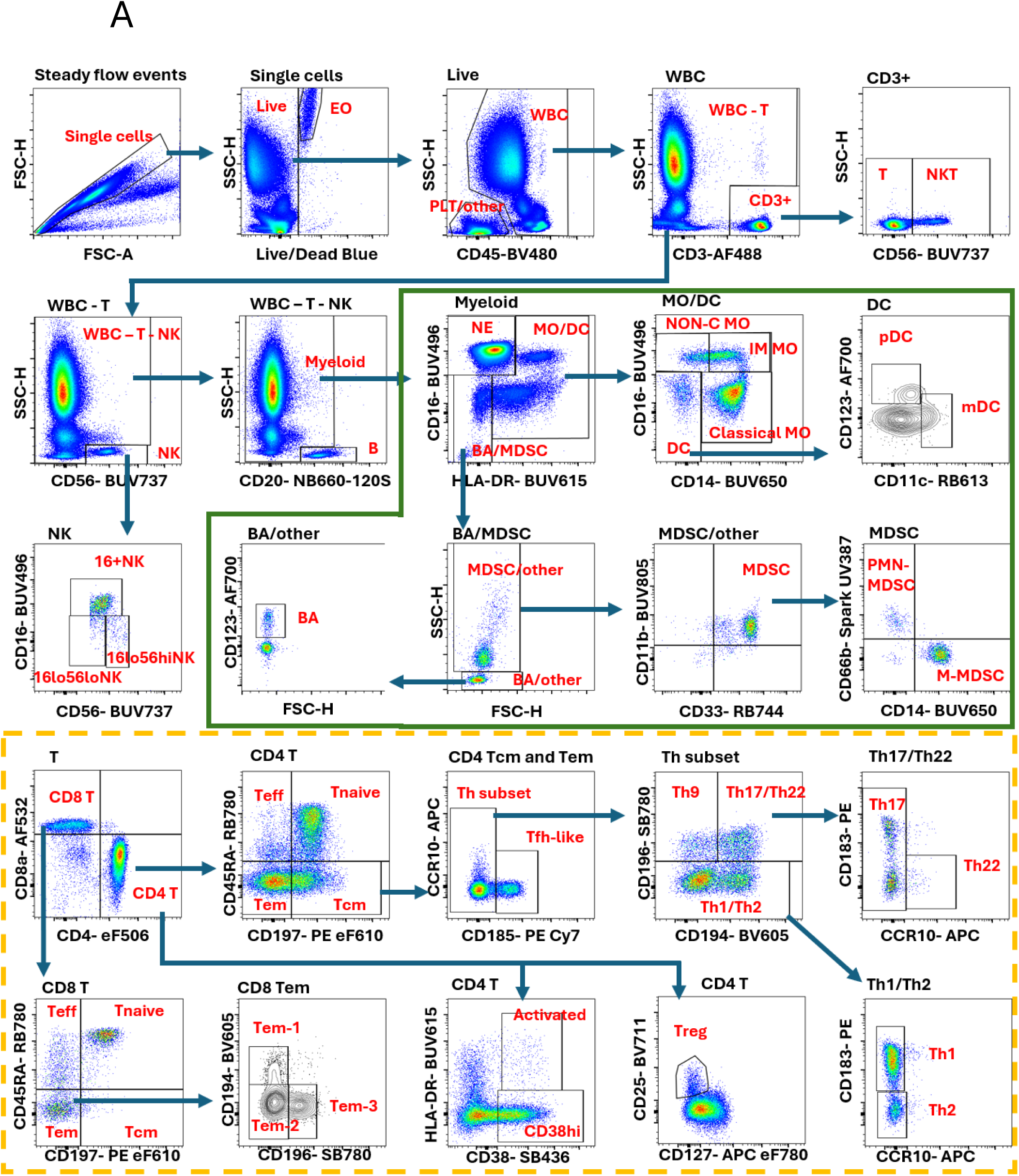
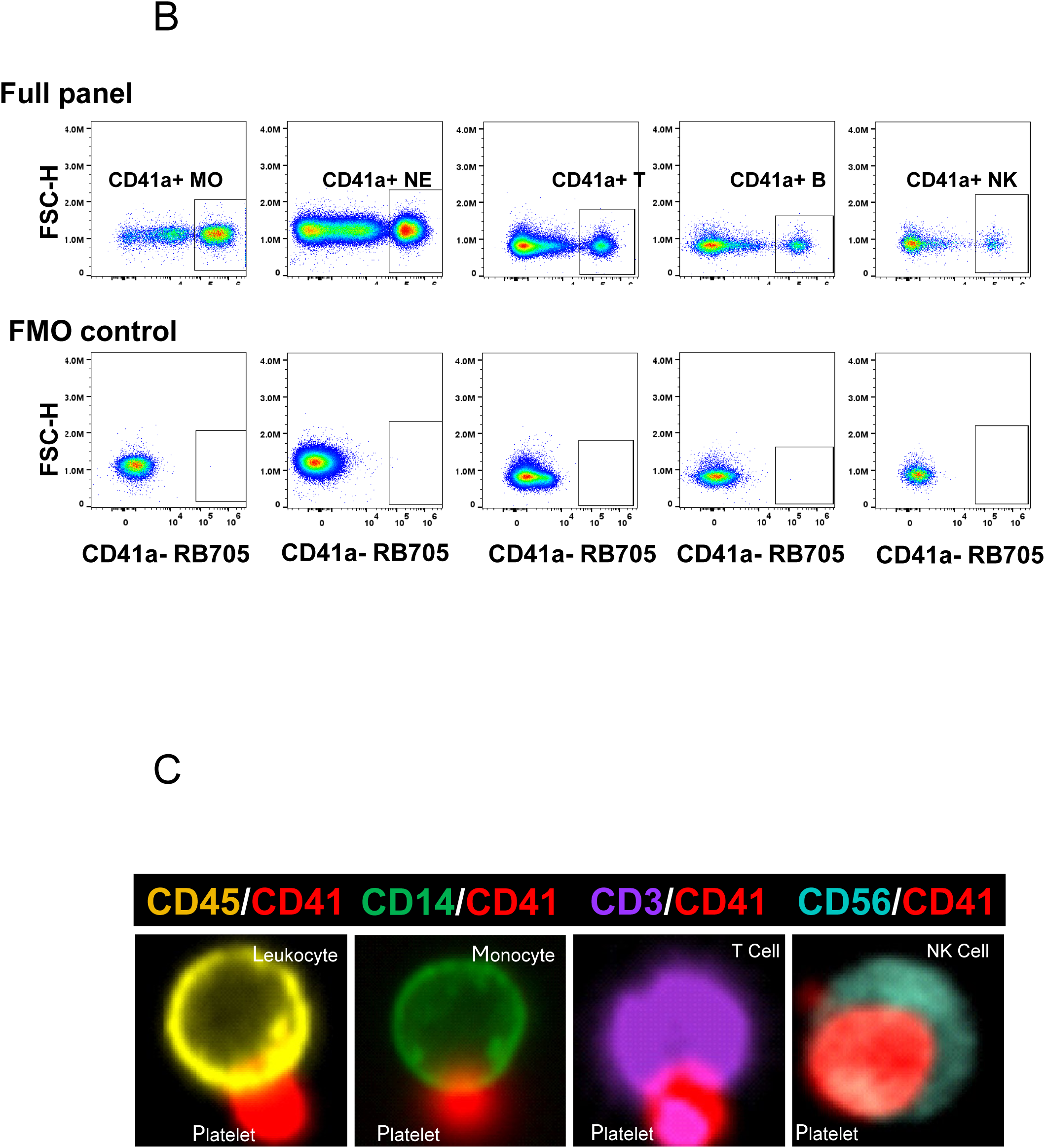
Gating strategy and PLA identification. (A) Representative spectral flow cytometry gating strategy for leukocyte populations. Bold headings denote the identified populations. Following identification of T cells, NK cells and B cells, the myeloid lineage gating strategy is shown within the solid-line box. Detailed T cell subset gating is shown in the dashed-line box. Contour plots were used for populations with low cell frequencies for better visualization. (B) Identification of PLAs for representative populations based on CD41 positivity (top panels). Fluorescence-minus-one (FMO) controls were used to demonstrate staining specificity. (C) Confirmation of CD41-positive leukocyte populations by imaging flow cytometry, validating the presence of platelet-associated leukocyte subsets identified by spectral flow cytometry, including monocytes, T cells and NK cells.

The relative abundance of each leukocyte subset was calculated by normalizing the population count to the total leukocyte (white blood cell (WBC)) count:

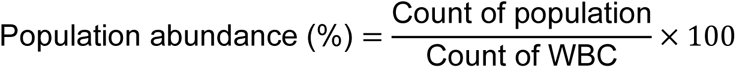

Absolute cell numbers were calculated using the corresponding WBC counts obtained from the CBC:

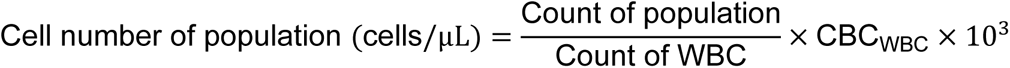

### UMAP analysis

Flow cytometry files (fcs files) were pre-processed (compensated, gated for single and viable cells and quality control checked using FlowAI). FlowSOM is an embedded algorithm within FlowJo for clustering cells with Self-Organizing Maps (SOMs), based on the behavior of chosen markers on all cells. We used FlowSOM to cluster cells by surface markers (including all markers in our panel excluding CD41). Clusters were then manually annotated and Uniform Manifold Approximation and Projections (UMAPs) were generated based on marker expression, limited to 150,000 cells per sample.

### Imaging flow cytometry

Whole blood samples from either ACD or sodium citrate tubes were centrifuged twice at 70 × g for 20 minutes at room temperature to remove platelet-rich plasma. The remaining cell fraction was diluted 1:1 with PBS and divided for pan-leukocyte and monocyte staining. For the pan-leukocyte panel, diluted samples were layered over Ficoll and centrifuged at 500 × g for 15 minutes at room temperature to enrich peripheral blood mononuclear cells. Cells were washed with PBS at 500 × g for 15 minutes, counted, and adjusted to 5 × 10^6^ cells/mL in PBS. Cells were then incubated with Fc receptor–blocking reagent for 10 minutes at room temperature in the dark, followed by staining with the antibody master mix for 20 minutes in the dark. After staining, samples were fixed using 4% paraformaldehyde. For the monocyte panel, PBS-diluted whole blood was stained directly after Fc receptor blocker with the indicated antibody master mix for 20 minutes at room temperature in the dark. After staining, samples were fixed using BD FACS Lysis buffer following manufacturer’s protocol. All fixed samples were then washed, resuspended in staining buffer, stored protected from light at 4°C, and acquired on the Amnis ImageStream X MKII imaging flow cytometer using 40x objective. IFC results were analyzed using Amnis IDEAS 6.2 software.

### Materials

All spectral flow antibodies can be found in Supplemental Table 1, all imaging flow antibodies can be found in Supplemental Table 2. Other protocol reagents can be found in Supplemental Table 3.

## Results

### 1. A Spectral Flow Cytometry Panel for Deep Immunophenotyping and PLA Classification

Built upon a widely adopted 21-color flow cytometry panel designed for comprehensive immunophenotyping of human peripheral blood (OMIP-042)^39,41^, we developed a 27-color (26 marker + Live/Dead Blue staining) spectral flow cytometry panel to simultaneously perform deep leukocyte immunophenotyping and detect PLAs. The complete list of antibodies is shown in Table 1 (Figure S1 shows the compensation matrix). We incorporated six additional markers: CD41, CD45, CD33, CD11b, CD66b, and CD15. CD41 was included as a platelet lineage marker to identify platelet-associated leukocytes, whereas CD45 was used to define total leukocytes. CD33 and CD11b were incorporated to improve identification of Myeloid-derived suppressor cells (MDSC). HLA-DR-/dimCD16-myeloid cells that are CD33+CD11b+ are defined as MDSCs.^42^ MDSC could be further divided into two major subsets: polymorphonuclear (PMN)-MDSC and monocytic (M)-MDSC by including CD66b. We also included CD15 to separate mature neutrophils (CD15+CD16+) and eosinophils (CD15+CD16-).

**Table 1.**
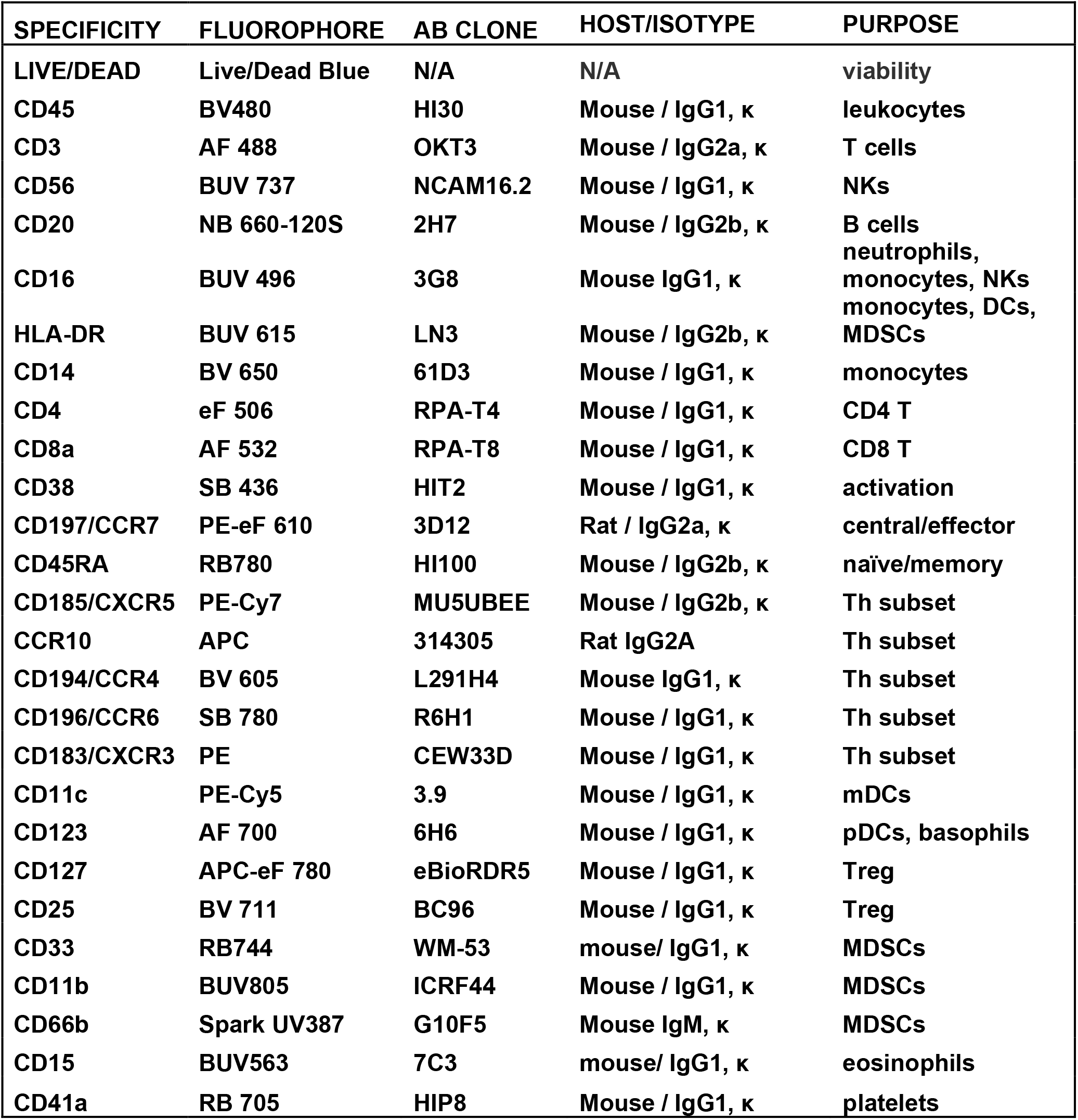
27-color panel for immunophenotyping and PLA detection.

The overall gating strategy is shown in Figure 1A, and the phenotypic definitions of leukocyte populations and subpopulations are summarized in Table 2. After exclusion of debris, doublets, and nonviable events, live CD45+ leukocytes (WBC) were identified. A subset of SSC^lo^ and CD45-contains free platelets (SSC^lo^CD45-CD41+). CD45+ leukocytes (white blood cell (WBC)) were then sequentially segregated into total T cells (SSC^lo^CD3+), natural killer (NK) cells (SSC^lo^CD56+), B cells (SSC^lo^CD20+) and myeloid cells (WBC-T-NK-B). NK cells further segregate into three populations according to CD56 and CD16 expression. Myeloid population contains both normal (neutrophils (NE), monocytes (MO), and dendritic cells (DC)) and atypical subsets (MDSCs). HLA-DR and CD16 were used to separate myeloid population into NE (HLA-DR-CD16+), MO/DC subset (HLA-DR+CD16+/-), and basophils (BA)/MDSC subset (HLA-DR-CD16-). Within MO/DC subset, CD14 and CD16 were used to identify various populations: DC (CD14-CD16-); classical MO (CD14^hi^CD16-); non-classical MO (CD14+CD16^hi^); and intermediate MO (CD14^hi^CD16+). Of the BA/MDSC subset, BA was identified (SSC^lo^ and CD123+); and MDSCs to be (SSC^hi^CD33+CD11b+). MDSCs were further differentiated into M-MDSCs (CD14+CD66b-) and PMN-MDSCs (CD14-CD66b+). Total T cells contain regular T cells (CD3+CD56-) and nature killer T (NKT) cells (CD3+CD56+). Immunophenotyping of conventional T cells basically followed the gating strategy of OMIP-042 (Figure 1A). CD4 and CD8 markers were first used to identify CD4+ T cells and CD8+ cells. Then a set of cell surface markers were used to identify T cell activation, T cell memory status, and all major Th subsets. Of note, HLA-DR and CD38 expression identifies T cell activation status within any subset.

**Table 2.**
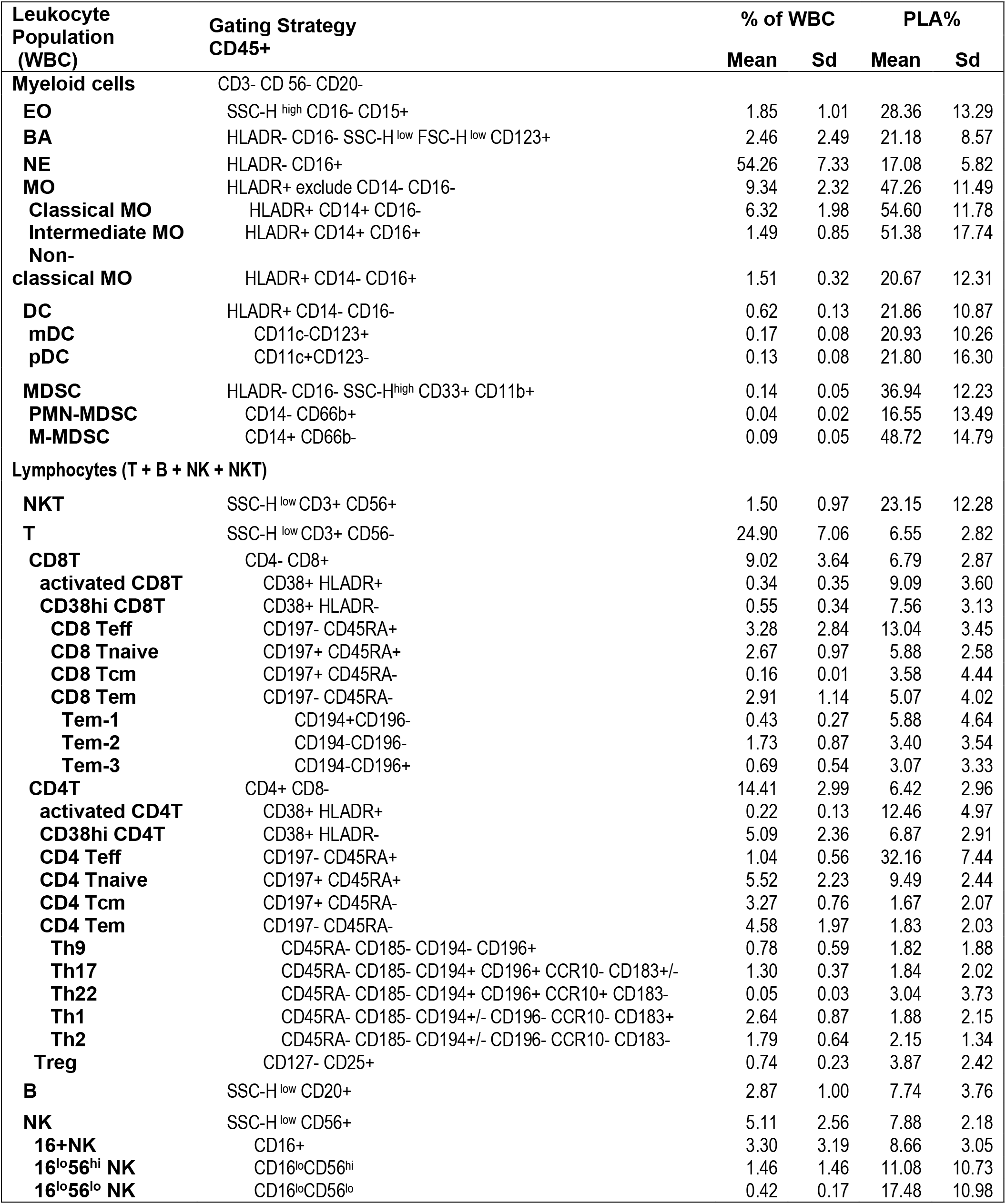
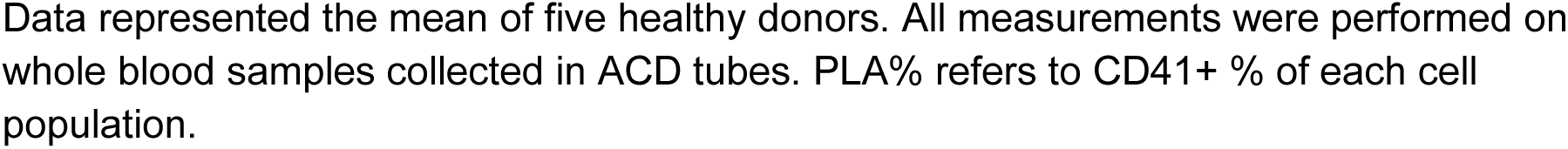
Immunophenotype of HC leukocytes and PLAs.

| Leukocyte Population (WBC) | Gating Strategy CD45+ | % of WBC |  | PLA% |  |
| --- | --- | --- | --- | --- | --- |
|  |  | Mean | Sd | Mean | Sd |
| **Myeloid cells** | CD3- CD 56- CD20- |  |  |  |  |
| **EO** | SSC-H<sup>high</sup> CD16- CD15+ | 1.85 | 1.01 | 28.36 | 13.29 |
| **BA** | HLADR- CD16- SSC-H<sup>low</sup> FSC-H<sup>low</sup> CD123+ | 2.46 | 2.49 | 21.18 | 8.57 |
| **NE** | HLADR- CD16+ | 54.26 | 7.33 | 17.08 | 5.82 |
| **MO** | HLADR+ exclude CD14- CD16- | 9.34 | 2.32 | 47.26 | 11.49 |
| **Classical MO** | HLADR+ CD14+ CD16- | 6.32 | 1.98 | 54.60 | 11.78 |
| **Intermediate MO** | HLADR+ CD14+ CD16+ | 1.49 | 0.85 | 51.38 | 17.74 |
| **Non-classical MO** | HLADR+ CD14- CD16+ | 1.51 | 0.32 | 20.67 | 12.31 |
| **DC** | HLADR+ CD14- CD16- | 0.62 | 0.13 | 21.86 | 10.87 |
| **mDC** | CD11c-CD123+ | 0.17 | 0.08 | 20.93 | 10.26 |
| **pDC** | CD11c+CD123- | 0.13 | 0.08 | 21.80 | 16.30 |
| **MDSC** | HLADR- CD16- SSC-H<sup>high</sup> CD33+ CD11b+ | 0.14 | 0.05 | 36.94 | 12.23 |
| **PMN-MDSC** | CD14- CD66b+ | 0.04 | 0.02 | 16.55 | 13.49 |
| **M-MDSC** | CD14+ CD66b- | 0.09 | 0.05 | 48.72 | 14.79 |
| **Lymphocytes (T + B + NK + NKT)** |  |  |  |  |  |
| **NKT** | SSC-H<sup>low</sup> CD3+ CD56+ | 1.50 | 0.97 | 23.15 | 12.28 |
| **T** | SSC-H<sup>low</sup> CD3+ CD56- | 24.90 | 7.06 | 6.55 | 2.82 |
| **CD8T** | CD4- CD8+ | 9.02 | 3.64 | 6.79 | 2.87 |
| **activated CD8T** | CD38+ HLADR+ | 0.34 | 0.35 | 9.09 | 3.60 |
| **CD38hi CD8T** | CD38+ HLADR- | 0.55 | 0.34 | 7.56 | 3.13 |
| **CD8 Teff** | CD197- CD45RA+ | 3.28 | 2.84 | 13.04 | 3.45 |
| **CD8 Tnaive** | CD197+ CD45RA+ | 2.67 | 0.97 | 5.88 | 2.58 |
| **CD8 Tcm** | CD197+ CD45RA- | 0.16 | 0.01 | 3.58 | 4.44 |
| **CD8 Tem** | CD197- CD45RA- | 2.91 | 1.14 | 5.07 | 4.02 |
| **Tem-1** | CD194+CD196- | 0.43 | 0.27 | 5.88 | 4.64 |
| **Tem-2** | CD194-CD196- | 1.73 | 0.87 | 3.40 | 3.54 |
| **Tem-3** | CD194-CD196+ | 0.69 | 0.54 | 3.07 | 3.33 |
| **CD4T** | CD4+ CD8- | 14.41 | 2.99 | 6.42 | 2.96 |
| **activated CD4T** | CD38+ HLADR+ | 0.22 | 0.13 | 12.46 | 4.97 |
| **CD38hi CD4T** | CD38+ HLADR- | 5.09 | 2.36 | 6.87 | 2.91 |
| **CD4 Teff** | CD197- CD45RA+ | 1.04 | 0.56 | 32.16 | 7.44 |
| **CD4 Tnaive** | CD197+ CD45RA+ | 5.52 | 2.23 | 9.49 | 2.44 |
| **CD4 Tcm** | CD197+ CD45RA- | 3.27 | 0.76 | 1.67 | 2.07 |
| **CD4 Tem** | CD197- CD45RA- | 4.58 | 1.97 | 1.83 | 2.03 |
| **Th9** | CD45RA- CD185- CD194- CD196+ | 0.78 | 0.59 | 1.82 | 1.88 |
| **Th17** | CD45RA- CD185- CD194+ CD196+ CCR10- CD183+/- | 1.30 | 0.37 | 1.84 | 2.02 |
| **Th22** | CD45RA- CD185- CD194+ CD196+ CCR10+ CD183- | 0.05 | 0.03 | 3.04 | 3.73 |
| **Th1** | CD45RA- CD185- CD194+/- CD196- CCR10- CD183+ | 2.64 | 0.87 | 1.88 | 2.15 |
| **Th2** | CD45RA- CD185- CD194+/- CD196- CCR10- CD183- | 1.79 | 0.64 | 2.15 | 1.34 |
| **Treg** | CD127- CD25+ | 0.74 | 0.23 | 3.87 | 2.42 |
| **B** | SSC-H<sup>low</sup> CD20+ | 2.87 | 1.00 | 7.74 | 3.76 |
| **NK** | SSC-H<sup>low</sup> CD56+ | 5.11 | 2.56 | 7.88 | 2.18 |
| **16+ NK** | CD16+ | 3.30 | 3.19 | 8.66 | 3.05 |
| **16<sup>lo</sup>56<sup>hi</sup> NK** | CD16<sup>lo</sup>CD56<sup>hi</sup> | 1.46 | 1.46 | 11.08 | 10.73 |
| **16<sup>lo</sup>56<sup>lo</sup> NK** | CD16<sup>lo</sup>CD56<sup>lo</sup> | 0.42 | 0.17 | 17.48 | 10.98 |

Data represented the mean of five healthy donors. All measurements were performed on whole blood samples collected in ACD tubes. PLA% refers to CD41+ % of each cell population.

CD41 positivity for each cell population was used to define PLAs for each cell population (Figure 1B). Imaging flow cytometry (IFC), which can visualize cell-cell interaction directly, further validated the PLAs observed (Figure 1C).

To ensure optimal performance, antibody titration was systematically conducted to determine optimal staining concentration for each antibody (Supplemental table 1, Figure S2A). Across tested CD41 antibody concentrations, PLA frequencies within major leukocyte populations remained stable (Figure S2B), supporting the specificity of the CD41-based gating strategy and indicating low background signal. Assay specificity was further evaluated using appropriate controls, including fluorescence minus one (FMO) controls and epitope-specific controls (Figure 1B).

As shown in Figure 2, our 27-color panel design enabled high-dimensional immunophenotyping of leukocyte populations. Uniform Manifold Approximation and Projection (UMAP) visualization of the total leukocytes (Figure 2A) and CD41+ leukocytes, which we defined as PLAs (Figure 2B) revealed a subset of CD41+ events within all major leukocyte populations. In table 2, we listed 40+ different leukocyte subpopulations and calculated the cell count and PLA% in each individual population (ACD-anticoagulated samples). To our knowledge, this is the first evidence of platelet associations with some of these populations, such as circulating Th9 and Th22 cells.

**Figure 2:**
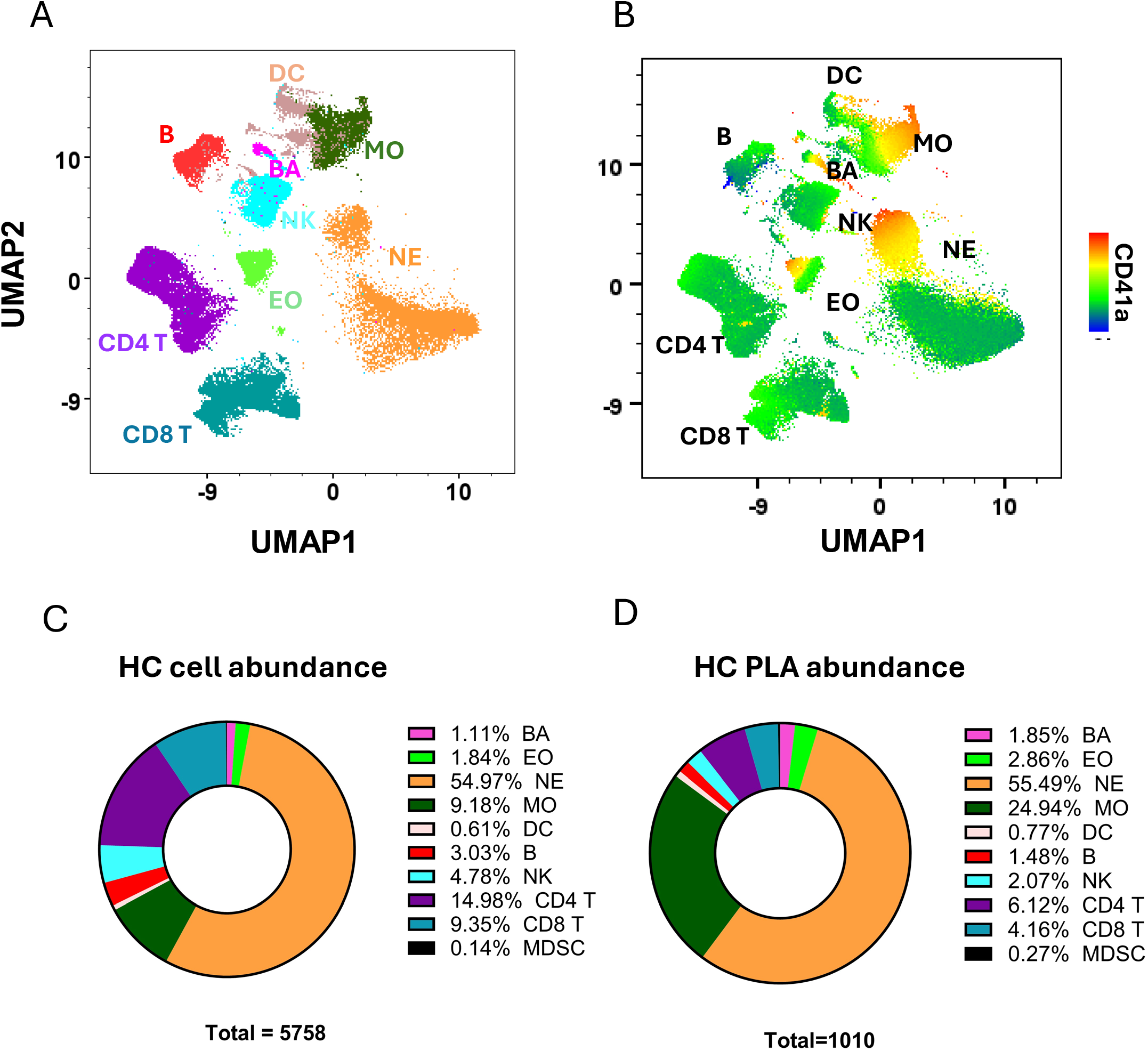
UMAP visualization of immune cell populations and PLA identification. (A-B) Representative UMAP visualization of major immunophenotyping (A) and a heatmap of CD41 expression (B) across the leukocytes from healthy donors. Monocytes (MO) and neutrophils (NE) showed highest CD41 expression. MO: monocytes, NE: neutrophils, BA: basophils, EO: eosinophils. (C) Distribution of major leukocyte subpopulations in white blood cells (WBC) from healthy donor whole blood collected in ACD (n=5). (D) Constitution of PLAs in the CD41+ WBC, showing neutrophils (orange) and monocytes (green) as the major platelet-binding populations. The cell count (cells/µL) shown on the bottom.

The frequency of platelet-leukocyte aggregates (PLA%) varied substantially among cell populations, ranging from less than 2% for CD4 memory T cells to greater than 50% for monocyte subsets (Table 2, Figure 2C-D). Myeloid populations, including monocytes (MO), neutrophils (NE), eosinophils (EO), and basophils (BA), exhibited markedly higher PLA% than lymphocyte populations, as indicated by both higher PLA% (Table 2) and higher CD41 MFI (Figure 2B). Although monocytes accounted for less than 10% of total leukocytes in healthy donors, they represented approximately 25% of total PLAs, indicating a disproportionate enrichment of platelet–monocyte aggregates (Figure 2D).

### 2. Reducing leukocyte-free platelet carryover and coincidence

Literature on PLA revealed significantly varied baselines in the healthy population, which may be due to technical variations, such as in anticoagulant choice, processing time, and sample preparations. We aim to optimize and standardize the PLA measurement protocol to reduce technical variations.

High throughput flow cytometry analysis enables analysis of millions of events per minute but also brings CD41+CD45+ coincidence when free platelets pass through the flow cytometer simultaneously that may be misclassified as PLAs. This issue cannot be fully resolved by forward- and side-scatter parameters alone; True PLAs may have scatter properties like single leukocytes, since platelets are much smaller than leukocytes. A slower flow rate for data acquisition improves signal resolution and reduces coincidence detection (Figure S3).

If the coincidence is attributed to leukocyte-free platelets, removing the leukocyte-free platelets prior to acquisition should reduce the coincidence rate. To examine this, we attempted to remove leukocyte-free platelets after the sample staining and fixation step. We examined the PLA% under 4 conditions: 50 × g for 10 minutes, 100 x g for 10 minutes, 200 × g for 10 minutes, and 400 x g for 5 minutes. We found that centrifugation at 50 × g for 10 minutes reduced free platelet carryover by >80% (Figure 3A) while maintaining majority (>80%) of total white blood cell counts. The 50 x g for 10 minutes also reduced PLA%, indicating reducing leukocyte-free platelets are important to reduce the noise of the assay (Figure S4).

**Figure 3:**
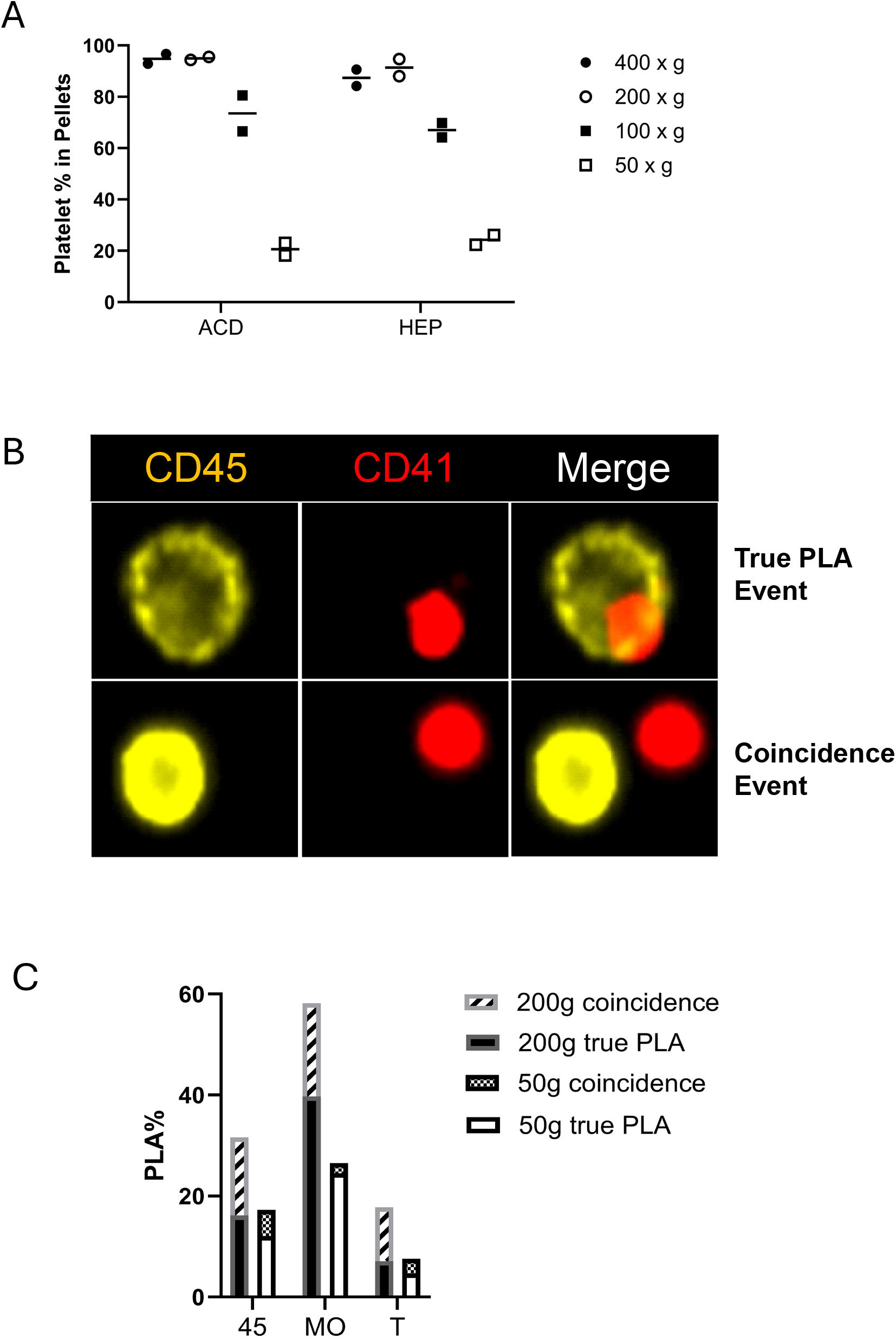
Reducing free platelets in stained samples helped to reduce coincidence events. (A) Centrifugation at 50 x g for 10 min removed majority of free platelets (>80%). Blood samples collected from healthy donors into either ACD or HEP were used. (B) Representative images of true PLA and a false PLA (coincidence event) by imaging flow cytometry. (C) The percentage of coincidence and true PLA in total leukocytes (CD45+), monocytes and T cells.

We further validated the effectiveness of the platelet removal step using IFC. IFC captures high resolution images of each cell. It enables separation of true PLA from coincidence events. Representative images of true positive and coincidence (false positive) events are shown in Figure 3B. As shown in Figure 3C, coincidence events were markedly reduced when a 50 × g centrifugation was used compared to a 200 × g speed.

### 3. Effect of anticoagulants

The choice of anticoagulants has been shown to significantly alter the overall PLA percentage.^25–27,30^ Whether different anticoagulants would affect different cell populations comparably or differentially, or what would be the preferred choice of anticoagulant regarding a specific cell population, remained unclear. We compared four commonly used anticoagulants: sodium citrate (CIT), acid citrate dextrose (ACD), EDTA, and heparin (HEP).

In healthy control samples, heparinized blood samples generally preserve lymphoid populations well, whereas EDTA samples showed a trend of modestly higher representation of myeloid populations (Figure 4A). Within each lineage, the distribution of subsets is highly comparable among different anticoagulants (data not shown). However, PLAs are more susceptible to anticoagulant choice, especially interactions between platelets and myeloid populations (PMA and PNA), whereas interactions between platelets-lymphocytes (T, B, and NK cells) are less affected (Figure 4B). This could be due to the status of platelets under different anticoagulants. For example, platelets are known to be activated when collected into HEP and subject to centrifugation.^43^ HEP samples followed by CIT samples showed higher platelet–monocyte aggregates and platelet-neutrophil aggregates compared with other anticoagulants. ACD samples showed lower PLA%. Interestingly, although EDTA are known to induce platelet swelling and activation, EDTA-treated samples generally showed the lowest PLA frequencies in monocytes and neutrophils.^44,45^

**Figure 4:**
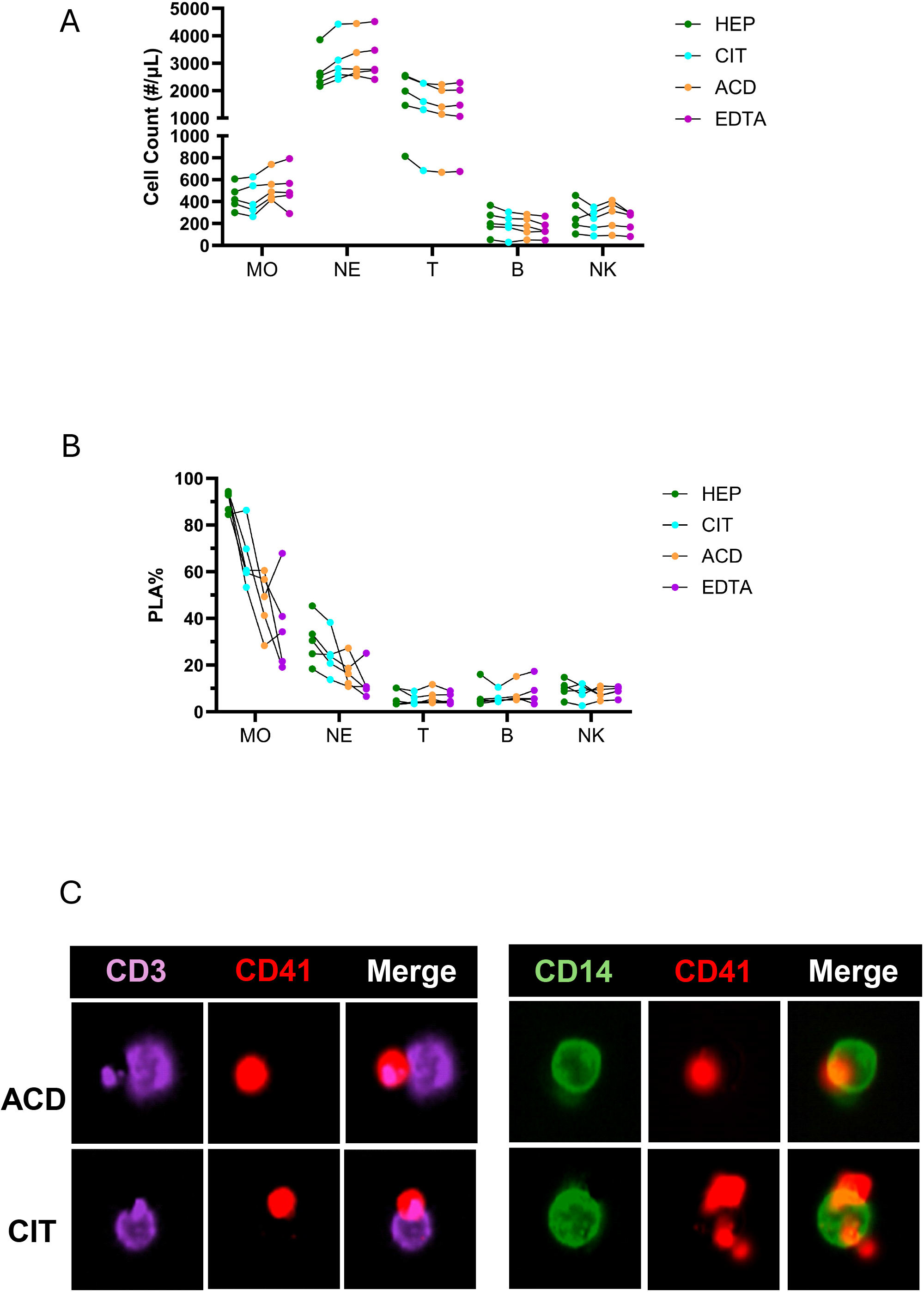
Effect of anticoagulants on immunophenotyping and PLA detection. Blood was drawn into 4 different anticoagulants in parallel for comparison. (A) Cell counts of major leukocyte populations. (B) CD41+ MO and CD41+ NE are sensitive to choice of anticoagulants. This represents 5 healthy donors examined. C, Imaging flow example of platelet aggregates in T cells (left) and monocytes (right). We consistently observed multiple platelets bound to monocytes in samples collected in sodium citrate (CIT) tubes.

Compared to HEP and EDTA, both CIT and ACD are known to better preserve the physiological status of platelets.^46,47^ To better understand the difference, we compared CIT and ACD samples using IFC. Monocytes collected in sodium citrate tubes often have multiple platelets bound together with distinct morphologic changes (platelet satellitism), compared to the rounded single platelet binding within the ACD tubes (Figure 4C), indicating better preservation of platelet morphology in ACD. Moreover, in PLA analysis, the separation of the CD41⁺ and CD41⁻ populations were more distinct in samples collected in ACD as compared to CIT tubes (Figure S5). We chose ACD for further studies in the patient cohort as discussed below.

### 4. Protocol Optimization and Assay Quality

In addition to anticoagulants, we further optimized the assay on sample processing, handling time, storage time, and RBC lysis methods.

We examined the effect of sample dilution on PLA measurements. As shown in Figure S6A, dilution of whole blood prior to staining resulted in a marked reduction in PLA% across major leukocyte populations. Increasing the volumes of fixation and RBC lysis reagents also led to a decrease in PLA%, although the effect was less pronounced (Figure S6B). Following fixation and storage, dilution of samples immediately before flow cytometric acquisition caused an additional decrease in the PLA% (Figure S6C). These findings indicate that PLA measurements are highly sensitive to sample dilution at multiple stages of the workflow. Dilution before fixation may alter the extracellular milieu and the stability of platelet–leukocyte interactions. Standardized sample handling procedures, including the volume of the assay system, is important for reproducibility when measuring PLAs.

We also examined the time from blood collection to sample processing. Consistent with studies by others, we found that samples processed within 2 hours of blood collection maintained consistent PLA measurements, whereas longer delays increased variability and/or altered PLA% (Figure 5A).^23,48^

**Figure 5:**
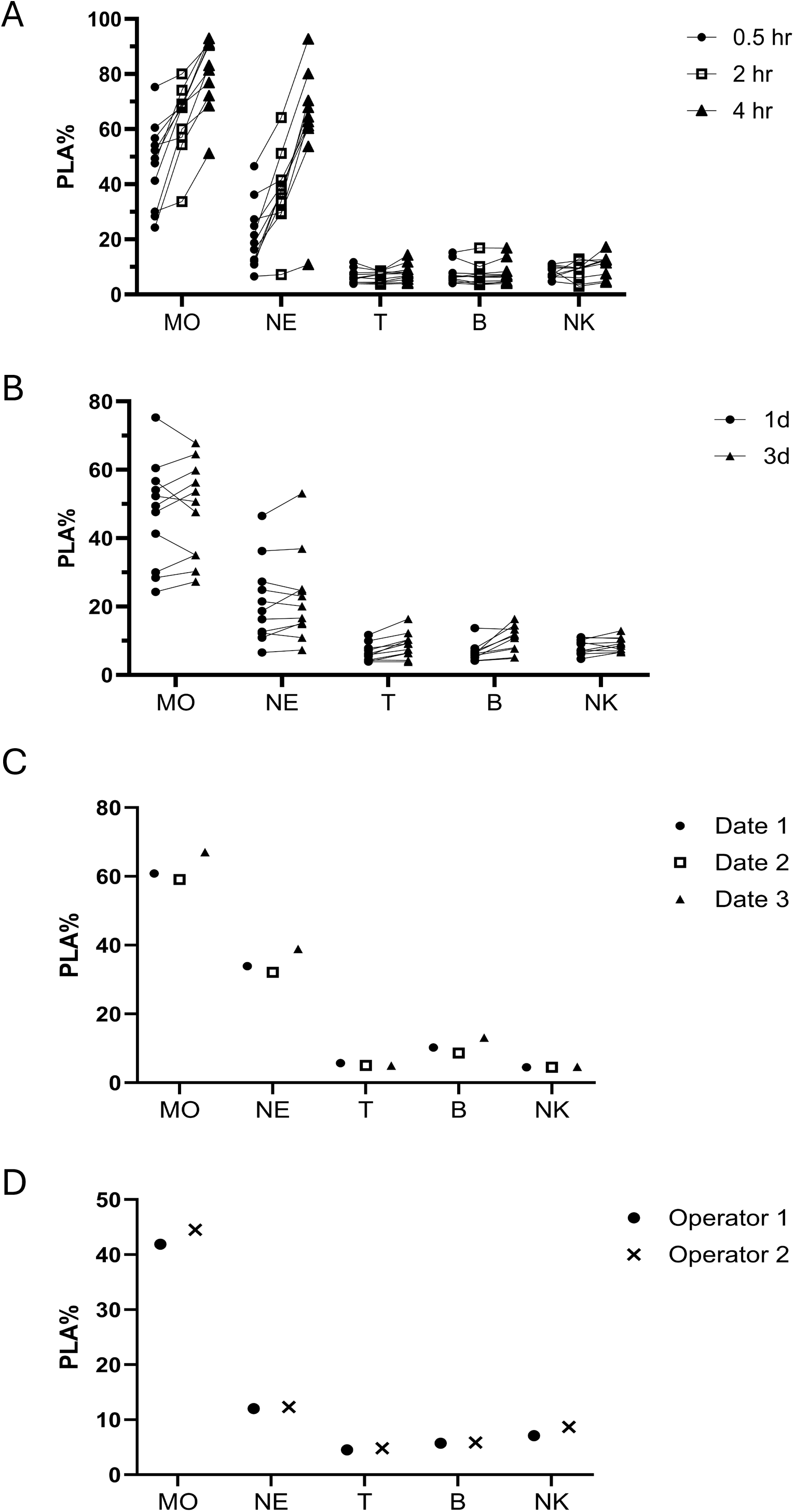
Assay stability and reproducibility. (A) Whole blood specimens were processed at 0.5, 2, and 4 h after collection to assess the effects of delayed processing on platelet–leukocyte aggregate percentages (PLA%). PLA% increased over time in monocytes and neutrophils but remained stable in lymphoid cell populations. (B) Following fixation, PLA% remained stable for at least 3 days during storage at 4°C, with minimal changes over the storage period. (C) PLA measurements from the same healthy donor on three separate blood collection days were highly comparable, supporting within-donor reproducibility. (D) PLA measurements obtained by different operators using blood from the same individual were highly consistent, supporting inter-operator reproducibility.

We also examined sample fixation and storage conditions. Fixation and RBC lysis combined or separately did not alter the results (data not shown). After fixation, samples stored at 4°C maintained stable fluorescence signals and PLA measurements for at least 3 days, providing flexibility for delayed flow cytometric acquisition after sample preparation (Figure 5B).

We next assessed the reproducibility and stability of our updated protocol. Duplicate samples prepared from the same blood draw showed minimal technical variability (Figure S7). Repeated blood draws from the same healthy donor across multiple days demonstrated low inter-day variation (Figure 5C), and samples processed by different operators showed minimal inter-operator variability (Figure 5D). These findings support the reproducibility of the assay across sample replicates, independent blood draws, and operators.

### 5. Finalized operating workflow

Based on the optimization and validation analyses described above, all subsequent experiments were performed using a standardized whole-blood workflow (Figure 6).

**Figure 6:**
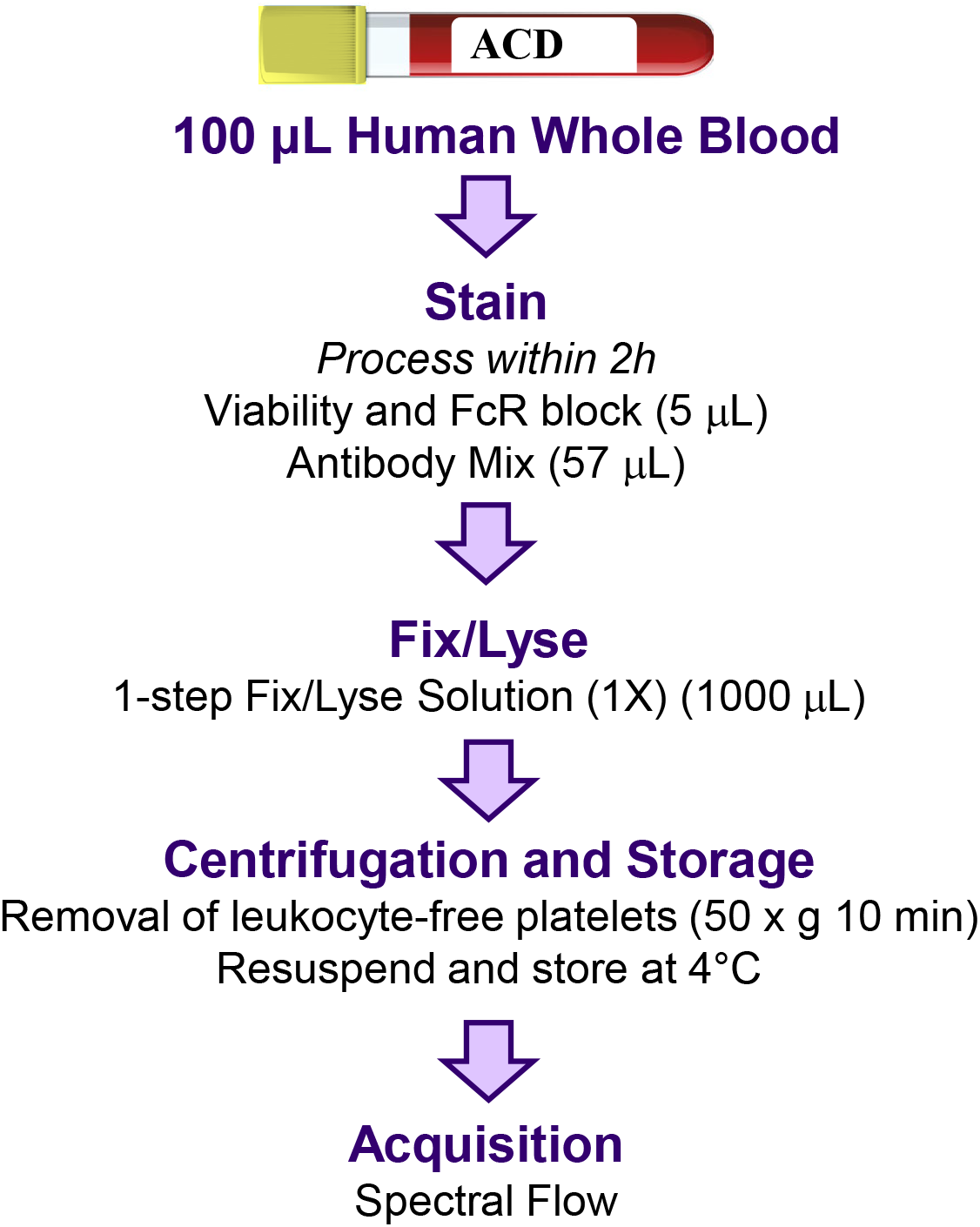
Optimized assay workflow. Briefly, 100 µL fresh whole blood was incubated with TruStain FcX (Biolegend) and Live/Dead Blue at room temperature for 10 min, then stained with antibody mixture (27-color panel) prepared with Brilliant Staining Buffer (BD Horizon) and Cellblox Blocking Buffer (Fisher) at room temperature, shaking for 30 min in dark. 1 mL 1xeBioscience™ 1-step Fix/Lyse Solution (Invitrogen) was then used to lyse red blood cells and fix stained samples at room temperature, shaking for 20 min in the dark. After centrifugation at 50 x g for 10 min, cell pellets were resuspended in 200 µL eBioscience™ Flow Cytometry Staining Buffer (Invitrogen) and stored at 4°C protected from light until acquisition.

### 6. Pilot study in sepsis and trauma patients

Having established the panel design, gating strategy, and optimized processing conditions, we next applied this assay to evaluate PLAs in two thromboinflammatory diseases of interest, sepsis and severe traumatic injury.

Compared with healthy controls (HC), both sepsis (SP) and trauma (TR) patients showed increased number and percentage of neutrophils (NE) within total leukocytes (Figure 7A, Figure S8A).^49^ Within the monocyte population, we observed increased number and percentage of M-MDSCs in both sepsis and trauma patients, consistent with studies by others (Fig 7A, Figure S8B), indicating the immunosuppression status of these patients.^50–53^ The PLA% within total neutrophils or M-MDSCs were largely unchanged compared to HC (Figure 7B). However, we observed consistent increase in number of platelet-binding neutrophils and M-MDSCs (Fig 7C, Figure S8B) in tandem with the pronounced neutrophilia and MDSC expansion (Figure 7A). In contrast, the number and percentage of non-classical monocytes within total monocytes decreased in both cohorts (Figure 7A, Figure S8B), so as the platelet-binding population (Figure 7C, Figure S8B).

**Figure 7:**
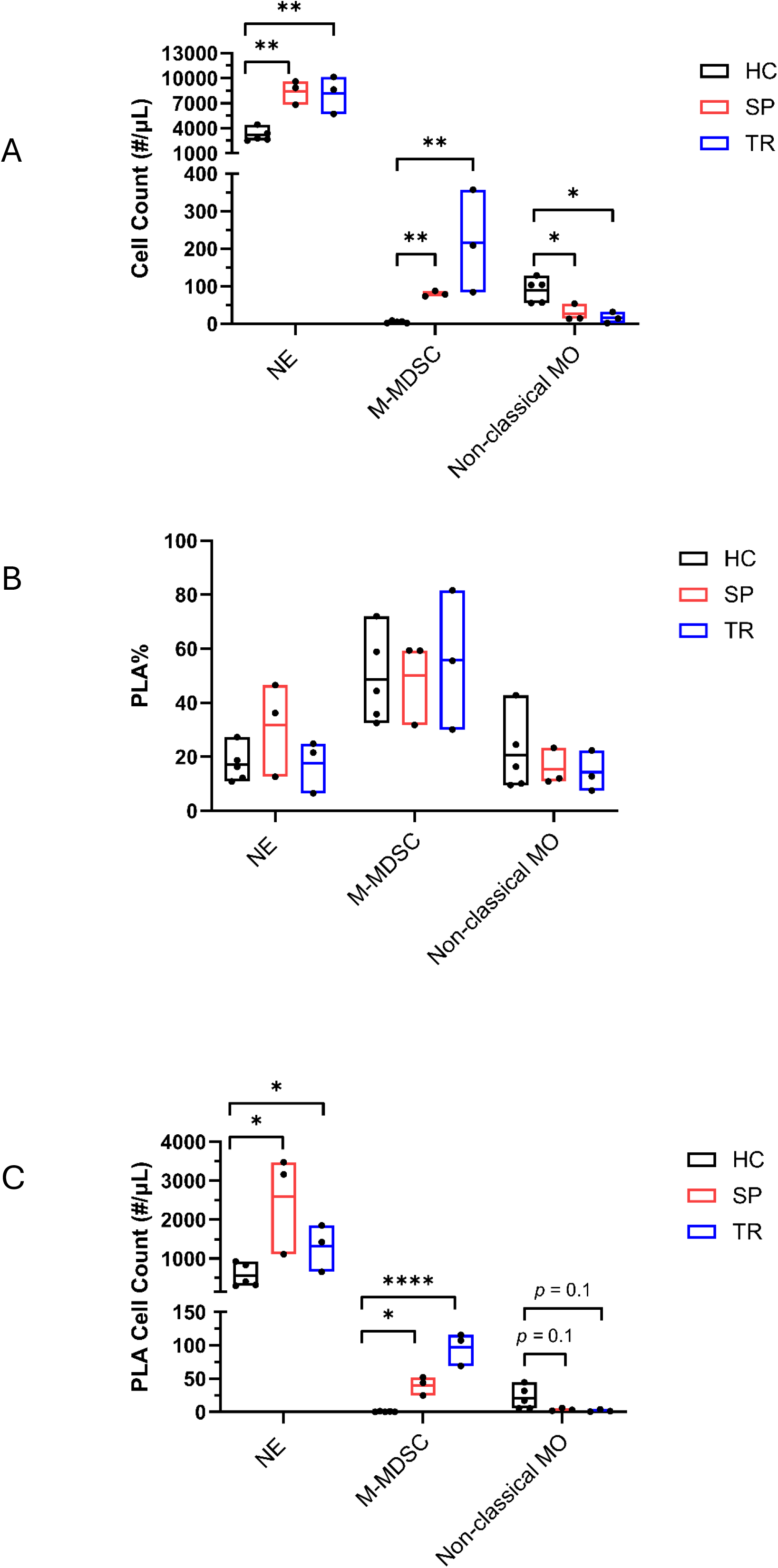
Deep immunophenotyping of patients with sepsis (SP) or trauma (TR). All statistics presented are ordinary one-way ANOVA Tukey’s multiple comparisons test, unless specified. (A) Significant differences in the absolute counts of myeloid cell subsets were observed among HC, SP, and TR groups. Both SP and TR patients exhibited markedly increased neutrophil (NE) and M-MDSC counts compared with HCs (NE: HC<>SP, *p* = 0.0028, HC<>TR, *p* = 0.0038; M-MDSC: HC<>SP, *p* = 0.0016 (Welch’s T Test), HC<>TR, *p* = 0.007), whereas non-classical monocyte counts were significantly reduced in both patient groups (Non-classical MO: HC<>SP, *p* = 0.0290, HC<>TR, *p* = 0.0127). Absolute cell counts (cells/µL) were calculated using complete blood count (CBC) data. (B) There was no significant change of PLA% associated with NE, M-MDSCs and Non-classical MO among HC, SP, and TR groups (NE: HC<>SP, *p* = 0.2203, HC<>TR, *p* = 0.9973; M-MDSC: HC<>SP, *p* = 0.9941, HC<>TR, *p* = 0.8723; Non-classical MO: HC<>SP, *p* = 0.7937, HC<>TR, *p* = 0.7136). (C) Despite similar PLA%, the absolute numbers of neutrophil-associated and M-MDSC-associated PLAs were significantly elevated in both SP and TR patients compared with HCs, reflecting the expansion of these myeloid populations in disease; on the other hand, significant decrease of non-classical MO resulted in associated PLA (NE: HC<>SP, *p* = 0. 0121(Unpaired T test), HC<>TR, *p* = 0.051 (Unpaired T test); M-MDSC: HC<>SP, *p* = 0.0127, HC<>TR, *p* = <0.0001; Non-classical MO: HC<>SP, *p* = 0.1409 (Unpaired T test), HC<>TR, *p* = 0.1115 (Unpaired T test)).

## Discussion

Platelet–leukocyte interactions are increasingly recognized as a core feature of thromboinflammation, yet their cellular distribution and biological interpretation remain largely unclear.^54,55^ Here we developed a 27-color whole-blood spectral flow cytometry protocol that enables comprehensive quantification of platelet association across >40 major myeloid and lymphoid compartments simultaneously, as well as deep immunophenotyping of these platelet-binding leukocytes within these major cell populations. Additionally, we demonstrated feasibility, reproducibility of known immune cell subset changes, and clinical relevance in blood samples from sepsis and severe trauma patients.^52^ For the broad application of PLA measurements, anticoagulant choice and handling procedures should be considered to minimize artifacts and variations.

Platelets are highly susceptible to ex-vivo activation, which can introduce artifacts into PLA measurement.^25,26,31,56^ Our protocol was designed to minimize sample manipulation and maintain the physiological state to accurately capture the level of PLA formation in peripheral blood. We observed that PLA measurement is sensitive to sample handling conditions, including the choice of anticoagulants, centrifugation, dilution factors, and preanalytical time. Our study underscores the importance of standardized processing protocols for accurate and reproducible assessment of PLAs. These observations extend earlier preanalytical work, showing that anticoagulant choice and delayed processing alters platelet-leukocyte aggregate measurement, and that different published protocols applied to the same sample yield markedly different results.^23^ Notably, we observed significant variations and artifacts from centrifugations that were not mentioned in previous studies, as detected by both spectral flow cytometry and IFC (Figure S4 and Figure 3)^22^

This comprehensive whole-blood assay allows quantitative comparison of platelet-leukocyte interactions across multiple leukocyte subsets, many of which were not reported before with direct interactions with platelets in the peripheral blood, such as Th9 and Th22 cells (Table 2). Future functional studies of platelet-binding Th9 and Th22 cells in healthy and disease status may be informative.

Our study revealed more than 5 fold differences in PLA% baseline comparing monocytes/neutrophils to T cells and B cells (Table 2). This indicates that the PLAs detected are not random artifacts, and platelets have preferred binding to myeloid cells. This may be attributed to the high expression of PSGL-1 or Mac-1 on myeloid cells, which have been shown able to bind platelet P-selectin and GPIb, or binding to fibrinogen in the plasma which can bind GPIIbIIIa on platelets. Furthermore, Mac-1-fibrinogen-Mac-1 binding could also help explain decreased PLA% upon blood sample dilution, as sample dilution decreases the concentration of fibrinogen.^57,58^ Although lymphocytes express PSGL-1 and integrins such as LFA-1, which can bind to ICAM-2 on platelets, the expression levels are relatively low on resting lymphocytes. It is also possible that platelet-lymphocyte interactions rely on other receptors.^59,60^

We purposefully designed our panel to better identify myeloid-derived suppressor cells (MDSCs) as compared to the OMIP-42 panel, of which cell populations play critical roles in the pathogenesis of both trauma and sepsis.^50–53^ In our current study, we successfully identified two major MDSC subsets: M-MDSCs and PMN-MDSCs, the immunosuppressive counterparts of monocytes and neutrophils, respectively. These subsets exhibited markedly different PLA frequencies. In healthy control samples, M-MDSCs displayed similar PLA% to conventional monocytes, whereas PMN-MDSCs showed similar PLA% to neutrophils (Table 2).^61–63^

The ability to simultaneously monitor multiple leukocyte subpopulations enabled direct comparison of these cellular responses and facilitated the identification of distinct cell type-specific effects that would likely have been overlooked using a more limited analytical approach. More broadly, these findings highlight the value of high-dimensional immunophenotyping, as simultaneous analysis of multiple leukocyte subsets enables the recognition of lineage-specific responses and reveals biological differences in platelet-leukocyte interactions that may not be apparent when examining cell populations individually.

In summary, we developed a robust and reliable spectral flow cytometry platform capable of simultaneously performing deep immunophenotyping and PLA measurements. Panels of comparable color number for whole-blood leukocyte immunophenotyping and for high-parameter platelet phenotyping have been published separately; The contribution here is their integration into a single, minimally manipulated whole-blood assay.^39,64,65^ Application of this platform revealed novel patterns of PLA formation and disease-associated alterations. However, given the limited sample size, these findings should be interpreted with caution, and it is premature to draw definitive conclusions regarding their biological and clinical significance. Ongoing studies are expanding patient enrollment and increasing statistical power to further validate these observations and assess their relevance in larger, independent cohorts.

## Acknowledgements

This work was supported by NIH/NIGMS grant R35GM142481 (S.B.), R35GM160360 (L.G.), and American Heart Association Award 938717 (L.G.), and University of Washington Institute of Translational Health Sciences NIAP Award (L.G.).

## Authorship

## Contribution

K.R.G., K.G.G., H.D., T.O., Y.T., and L.G. performed experiments; H.D., K.R.G., K.G.G., A.H., E.L.H., and E.Z. analyzed results; H.D., K.R.G., and K.G.G. generated figures; S.C.B., H.D., and L.G. designed the research and H.D., K.R.G., K.G.G., T.O., L.G., and S.C.B. prepared and finalized the manuscript.

