## Supplemental Tables for "Whole Blood Deep Immunophenotyping and Platelet-Leukocyte Aggregate Quantification by Spectral Flow Cytometry"

Supplemental Table 1.

| Description | Manufacturer | Catalog Number | Titration |
| --- | --- | --- | --- |
| Antibodies: Spectral Flow | | | |
| CD3 AF488 | Invitrogen | 53003742 | 1:800 |
| CD16 BUV496 | BD Biosciences | 612944 | 1:800 |
| CD56 BUV737 | Invitrogen | 367056642 | 1:400 |
| CD38 SB436 | Invitrogen | 62038942 | 1:400 |
| CD4 eFluro506 | Invitrogen | 69004942 | 1:400 |
| CD196 SB780 | Invitrogen | 78196942 | 1:200 |
| CD45RA RB780 | BD Biosciences | 755990 | 1:1600 |
| CD197 PE-eF610 | Invitrogen | 61197942 | 1:800 |
| CD185 PE-Cy7 | Invitrogen | 25918542 | 1:800 |
| CD25 BV711 | Invitrogen | 407025942 | 1:400 |
| CD183 PE | Invitrogen | 12183942 | 1:400 |
| CD11c RB613 | BD Horizon | 571110 | 1:1600 |
| CD123 AF700 | Invitrogen | 56123942 | 1:400 |
| CD127 APC eF780 | Invitrogen | 47127842 | 1:400 |
| CD194 BV605 | Biolegend | 359418 | 1:200 |
| CCR10 APC | R&D Systems | FAB3478A | 1:200 |
| CD8a AF532 | Invitrogen | 58008842 | 1:400 |
| CD14 BV650 | Invitrogen | 416014942 | 1:800 |
| HLADR BUV615 | Invitrogen | 366995642 | 1:800 |
| Live/Dead Blue Cell Stain | Invitrogen | L34961 | 1:1200 |
| CD11b BUV805 | Invitrogen | 368011842 | 1:800 |
| CD45 BV480 | Invitrogen | 414045942 | 1:400 |
| CD20 NB660-120S | Invitrogen | H076T03B08 | 1:800 |
| CD33 RB744 | BD OptiBuild | 758127 | 1:800 |
| CD41a RB705 | BD OptiBuild | 757711 | 1:400 |
| CD66b SparkUV387 | Biolegend | 305128 | 1:200 |
| CD15 BUV563 | BD Horizon | 568918 | 1:1600 |

Supplemental Table 2.

| Description | Manufacturer | Catalog Number | Chanel |
| --- | --- | --- | --- |
| Antibodies: Imaging Flow | | | |
| Leukocyte | | | |
| CD3 APC-eFluor 780 | ThermoFisher | 47-0037-42 | 12 |
| CD8 PE-Cy5.5 | ThermoFisher | 45-0088-42 | 5 |
| CD4 PE | ThermoFisher | 12-0049-42 | 3 |
| CD45 RA Super Bright 600 | ThermoFisher | 63-0458-42 | 10 |
| CD41a R718 | BD | 568058 | 11 |
| CD197 FITC | ThermoFisher | 11-1979-42 | 2 |
| CD62L PC7 | ThermoFisher | 25-0629-42 | 6 |
| Viability L/D Aqua | ThermoFisher | L34957 | 8 |
| NKT | | | |
| CD45 PE | BD | 555483 | 3 |
| CD8 PE-Cy5.5 | ThermoFisher | 45-0088-42 | 5 |
| CD3 APC-eFluor 780 | ThermoFisher | 47-0037-42 | 12 |
| CD41a R718 | BD | 568058 | 11 |
| CD56 FITC | ThermoFisher | 11-0566-42 | 2 |
| Viability L/D Aqua | ThermoFisher | L34957 | 8 |
| Monocyte | | | |
| CD14 PE-Cy5.5 | ThermoFisher | MHCD1418 | 5 |
| CD16 APC-eFluor 780 | ThermoFisher | 47-0168-42 | 12 |
| HLA-DR FITC | ThermoFisher | 11-9956-42 | 2 |
| CD41a R718 | BD | 568058 | 11 |
| CD11b PE | ThermoFisher | 12-0118-42 | 3 |
| CD33 e450 | ThermoFisher | 48-0337-42 | 7 |
| Viability L/D Aqua | ThermoFisher | L34957 | 8 |

Supplemental Table 3.

| Description | Manufacturer | Catalog number |
| --- | --- | --- |
| Supplies: General | | |
| Heparin blood collection tubes | BD | 367878 |
| EDTA blood collection tubes | BD | 367841 |
| ACD blood collection tubes | BD | 364606 |
| NaCitrate blood collection tubes | BD | 363083 |
| Supplies: Spectral Flow | | |
| Human TruStain FcX | Biolegend | 422302 |
| Polypropylene Test Tubes with Cap | Falcon | 352063 |
| Brilliant Stain Buffer | Invitrogen | 00440975 |
| CellBlox Plus Blocking Buffer | Invitrogen | C001T03F01 |
| Flow Cytometry Staining Buffer | Invitrogen | 00422226 |
| 1-step Fix/Lyse Solution (10X) | Invitrogen | 00533357 |
| Supplies: Imaging Flow | | |
| FcR Binding Inhibitor | Thermofisher | 14-9161-73 |
| FACS Lysing Solution | BD | 349202 |
| Ficoll-Paque | Cytiva | 17144002 |
| Paraformaldehyde | EMS | 15713-S |
