## Supplemental Figures for "Whole Blood Deep Immunophenotyping and Platelet-Leukocyte Aggregate Quantification by Spectral Flow Cytometry"

### Slide 1
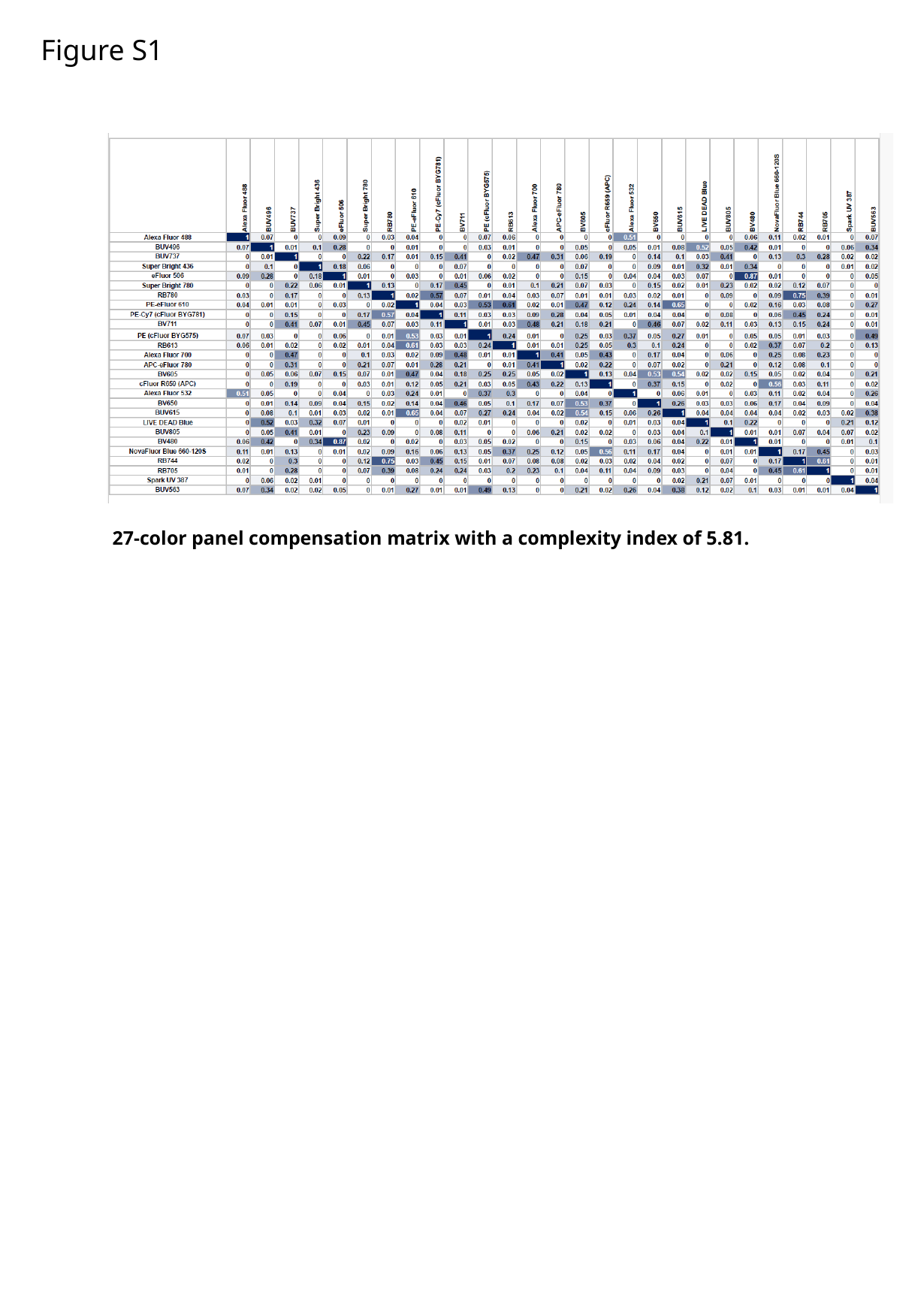

Figure S1
27-color panel compensation matrix with a complexity index of 5.81.

### Slide 2
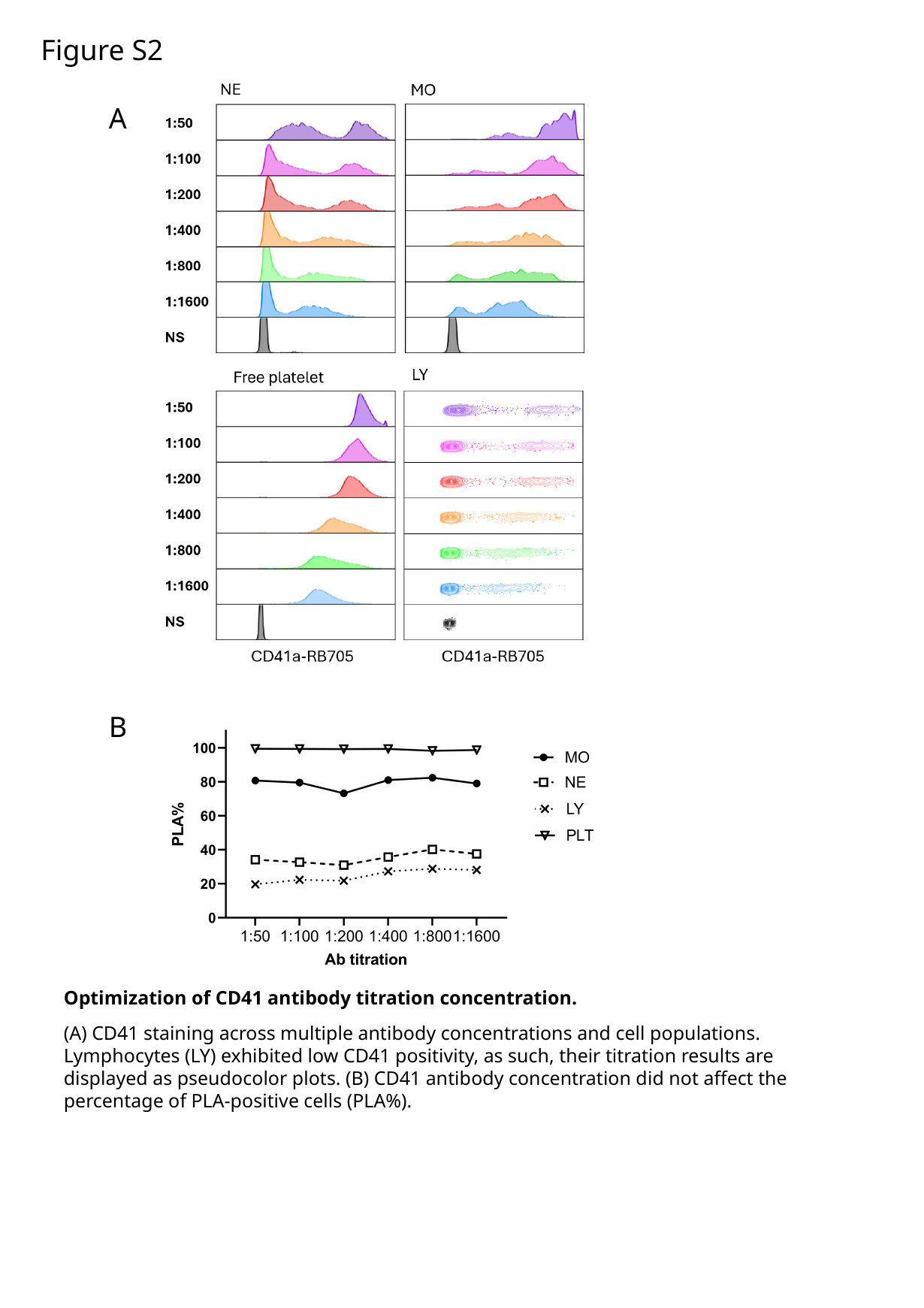

Figure S2
A
B
Optimization of CD41 antibody titration concentration.
(A) CD41 staining across multiple antibody concentrations and cell populations. Lymphocytes (LY) exhibited low CD41 positivity, as such, their titration results are displayed as pseudocolor plots. (B) CD41 antibody concentration did not affect the percentage of PLA-positive cells (PLA%).

### Slide 3
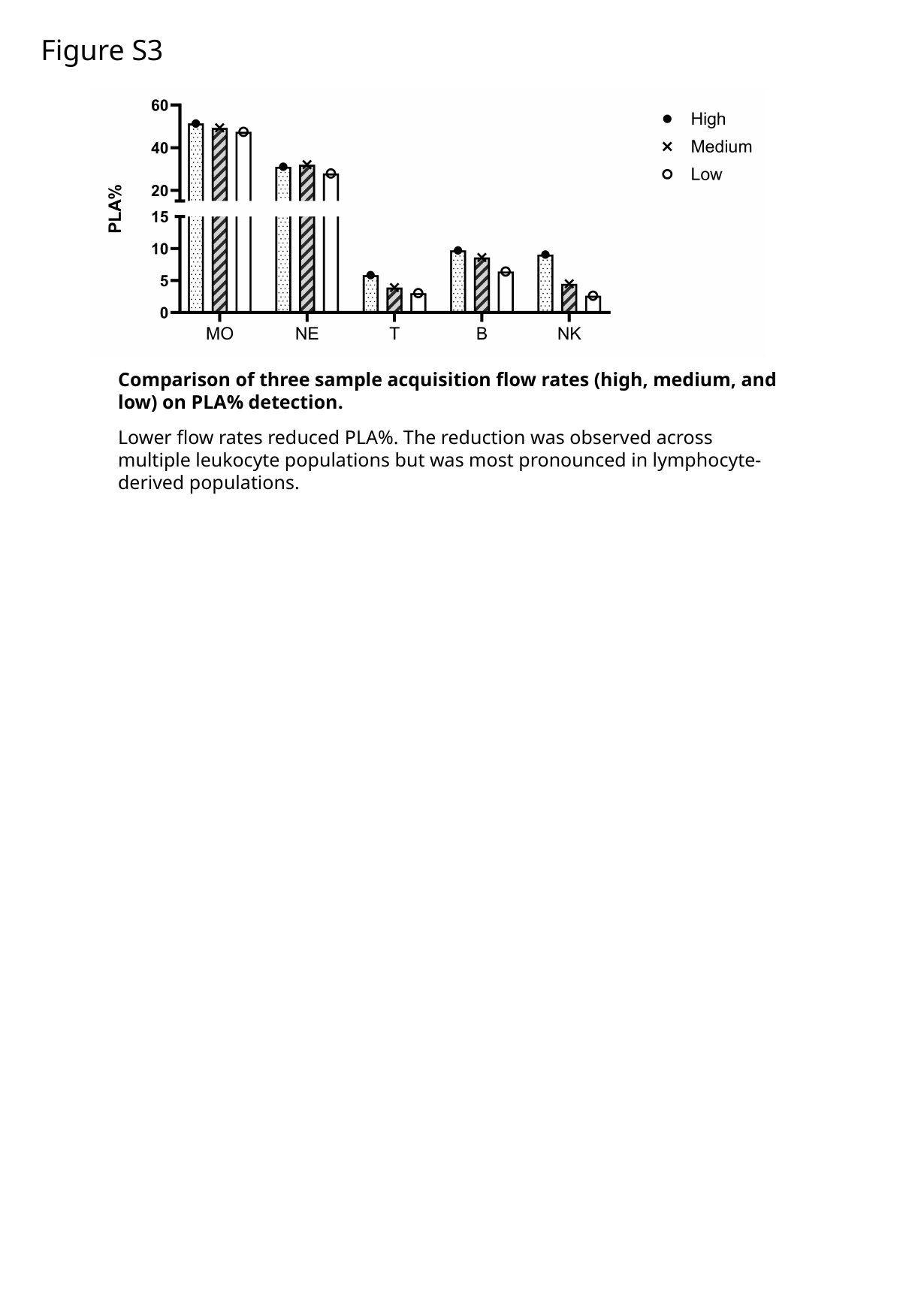

Figure S3
Comparison of three sample acquisition flow rates (high, medium, and low) on PLA% detection.
Lower flow rates reduced PLA%. The reduction was observed across multiple leukocyte populations but was most pronounced in lymphocyte-derived populations.

### Slide 4
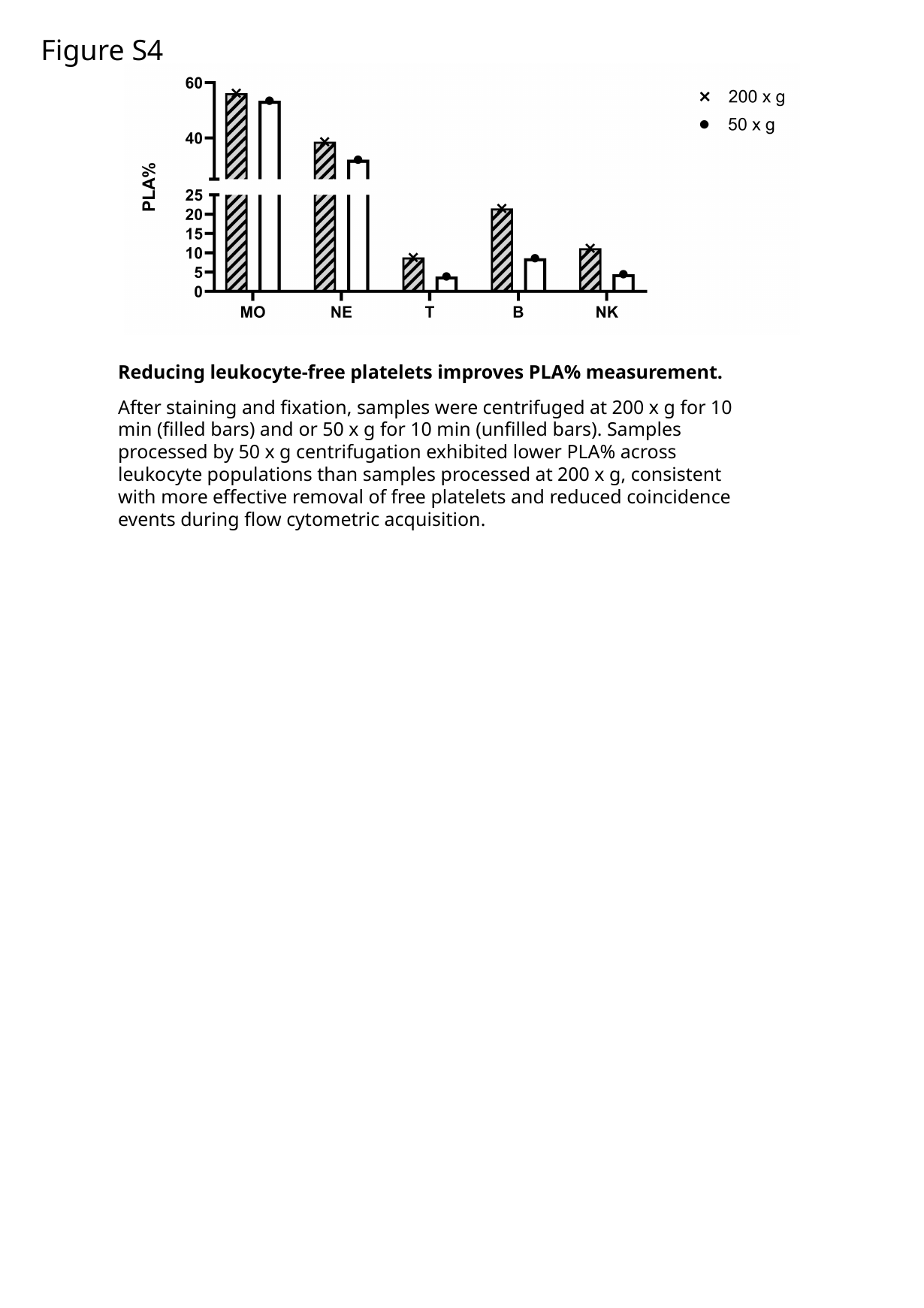

Figure S4
Reducing leukocyte-free platelets improves PLA% measurement.
After staining and fixation, samples were centrifuged at 200 x g for 10 min (filled bars) and or 50 x g for 10 min (unfilled bars). Samples processed by 50 x g centrifugation exhibited lower PLA% across leukocyte populations than samples processed at 200 x g, consistent with more effective removal of free platelets and reduced coincidence events during flow cytometric acquisition.

### Slide 5
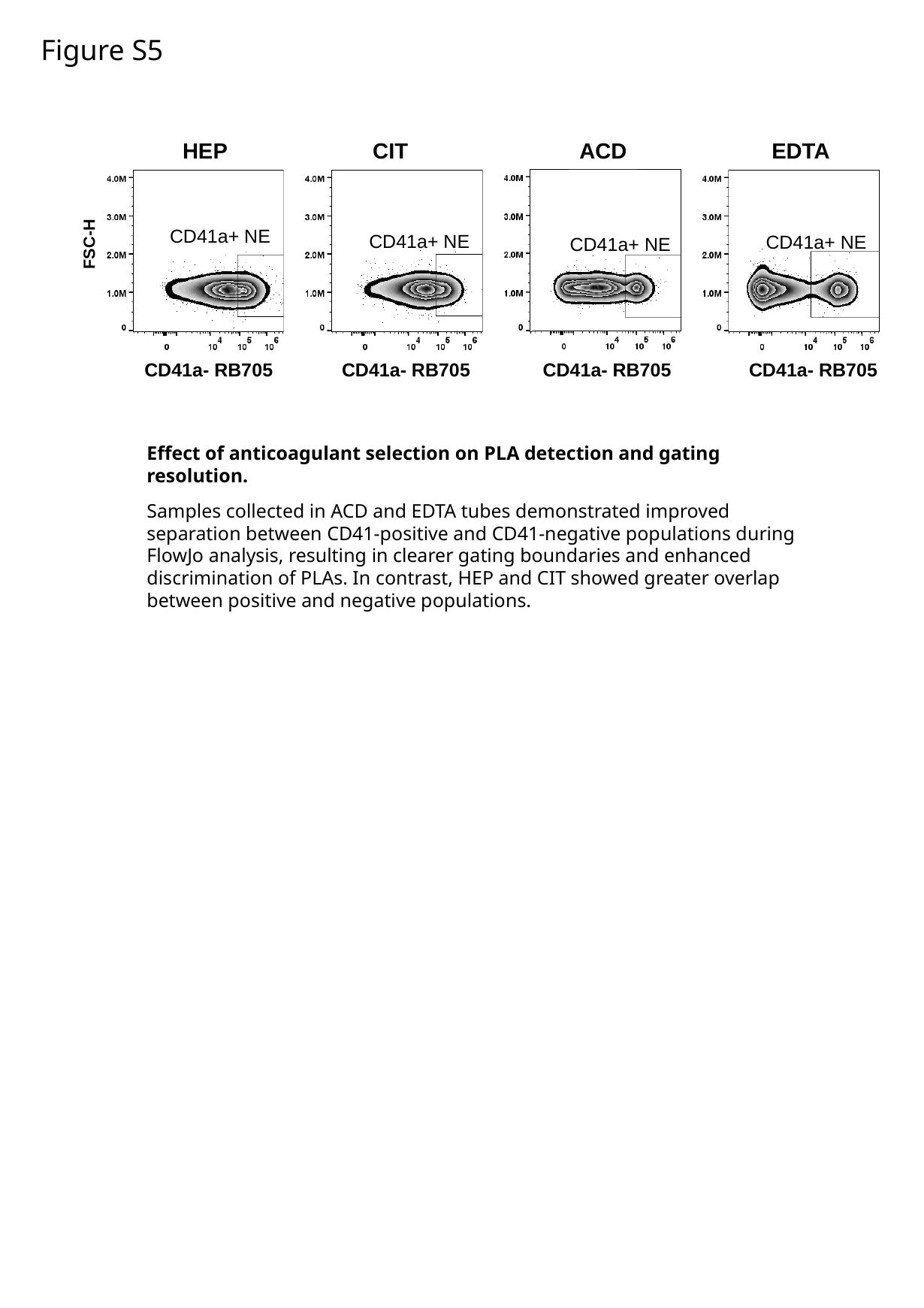

Figure S5
CIT
CD41a+ NE
CD41a- RB705
HEP
CD41a+ NE
CD41a- RB705
ACD
CD41a+ NE
CD41a- RB705
EDTA
CD41a+ NE
CD41a- RB705
FSC-H
Effect of anticoagulant selection on PLA detection and gating resolution.
Samples collected in ACD and EDTA tubes demonstrated improved separation between CD41-positive and CD41-negative populations during FlowJo analysis, resulting in clearer gating boundaries and enhanced discrimination of PLAs. In contrast, HEP and CIT showed greater overlap between positive and negative populations.

### Slide 6
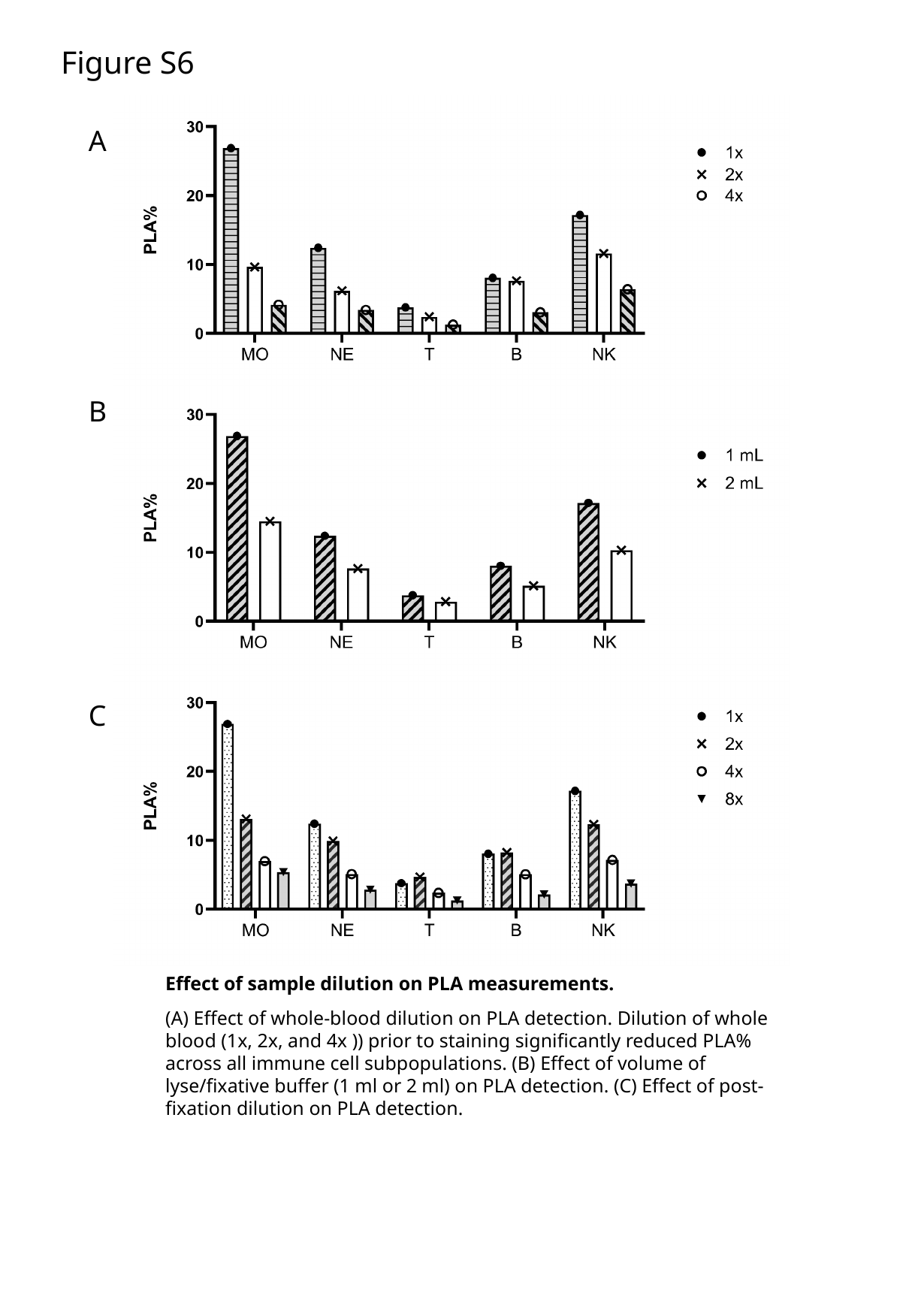

Figure S6
A
B
C
Effect of sample dilution on PLA measurements.
(A) Effect of whole-blood dilution on PLA detection. Dilution of whole blood (1x, 2x, and 4x )) prior to staining significantly reduced PLA% across all immune cell subpopulations. (B) Effect of volume of lyse/fixative buffer (1 ml or 2 ml) on PLA detection. (C) Effect of post-fixation dilution on PLA detection.

### Slide 7
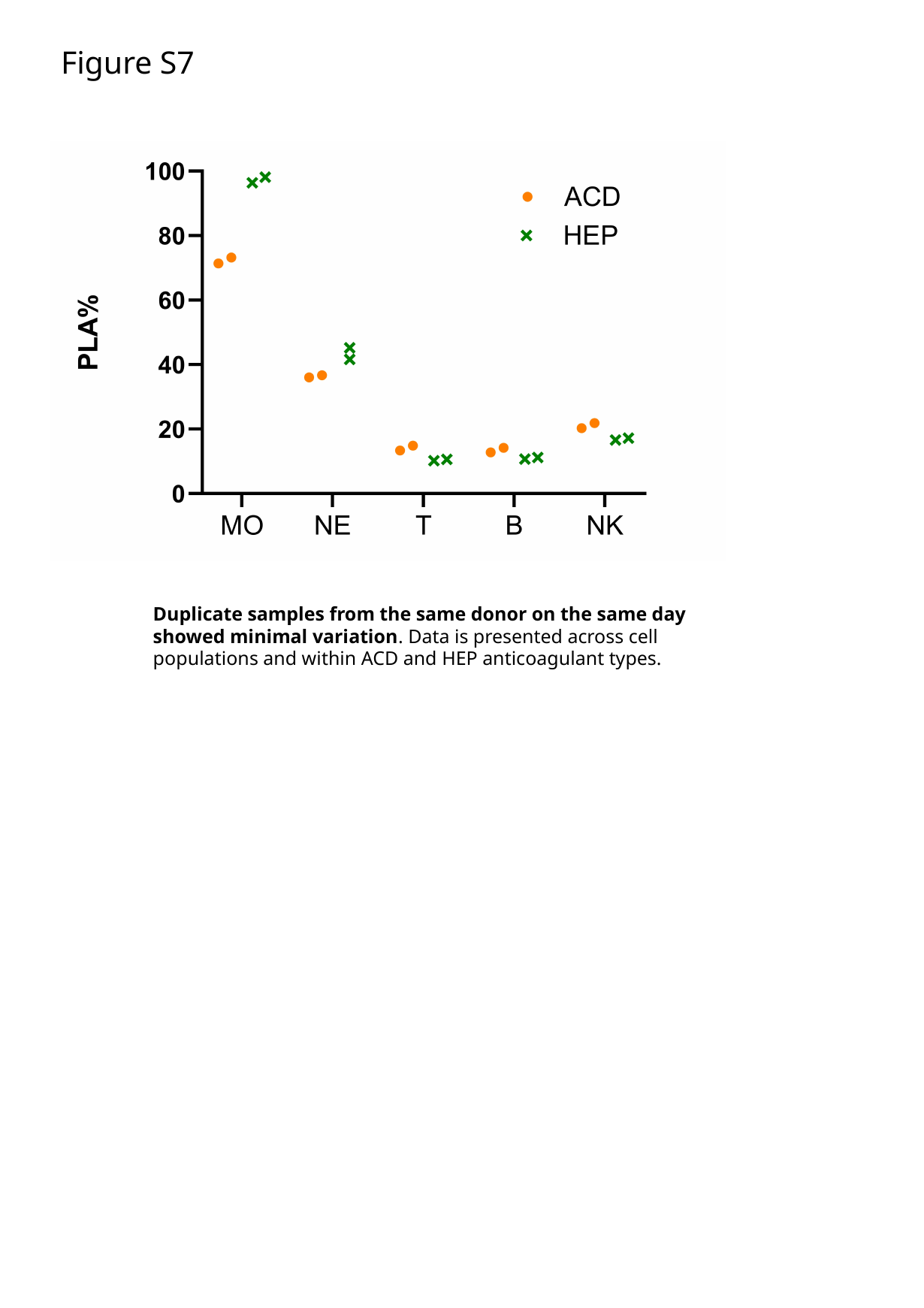

Figure S7
Duplicate samples from the same donor on the same day showed minimal variation. Data is presented across cell populations and within ACD and HEP anticoagulant types.

### Slide 8
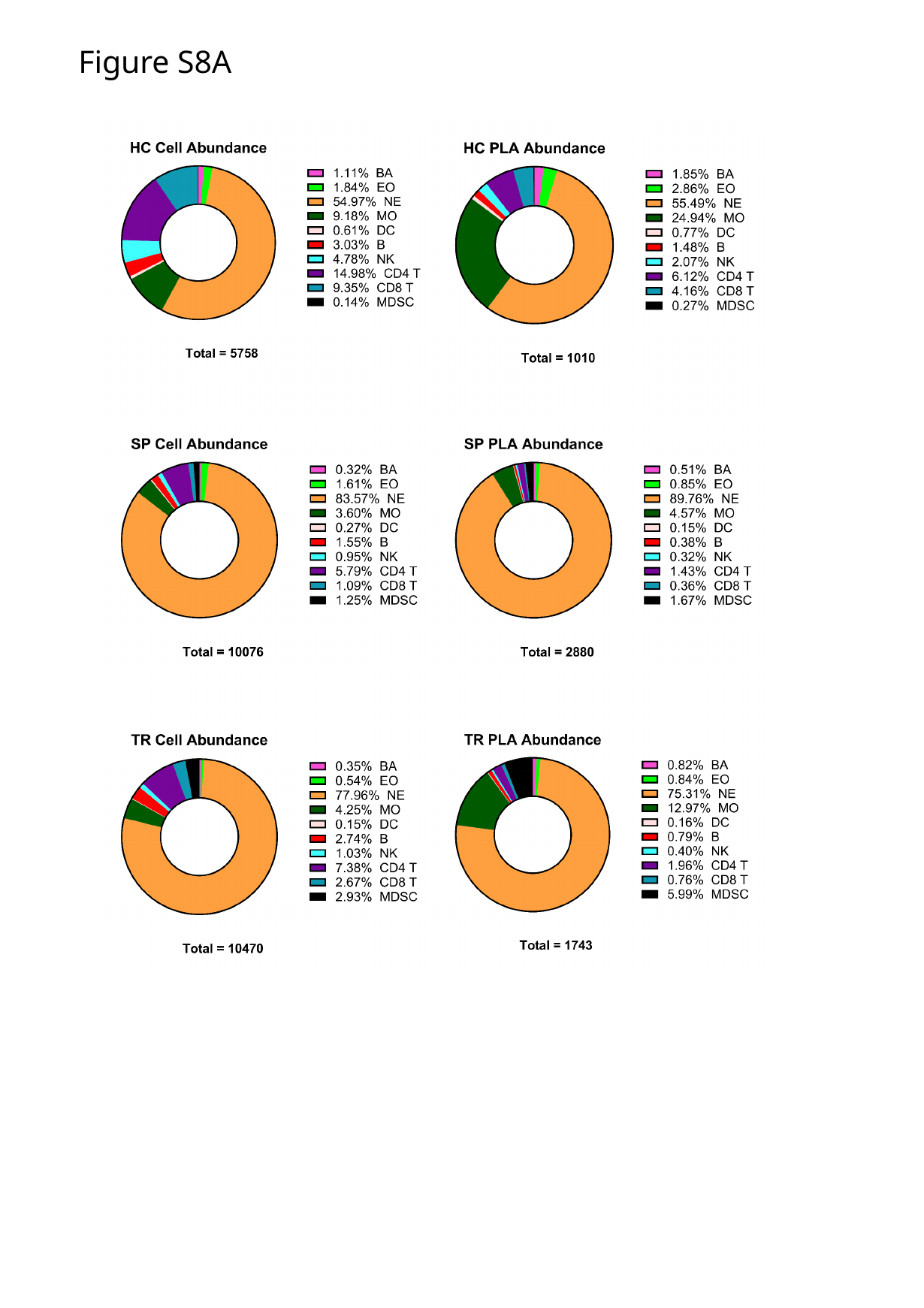

Figure S8A

### Slide 9
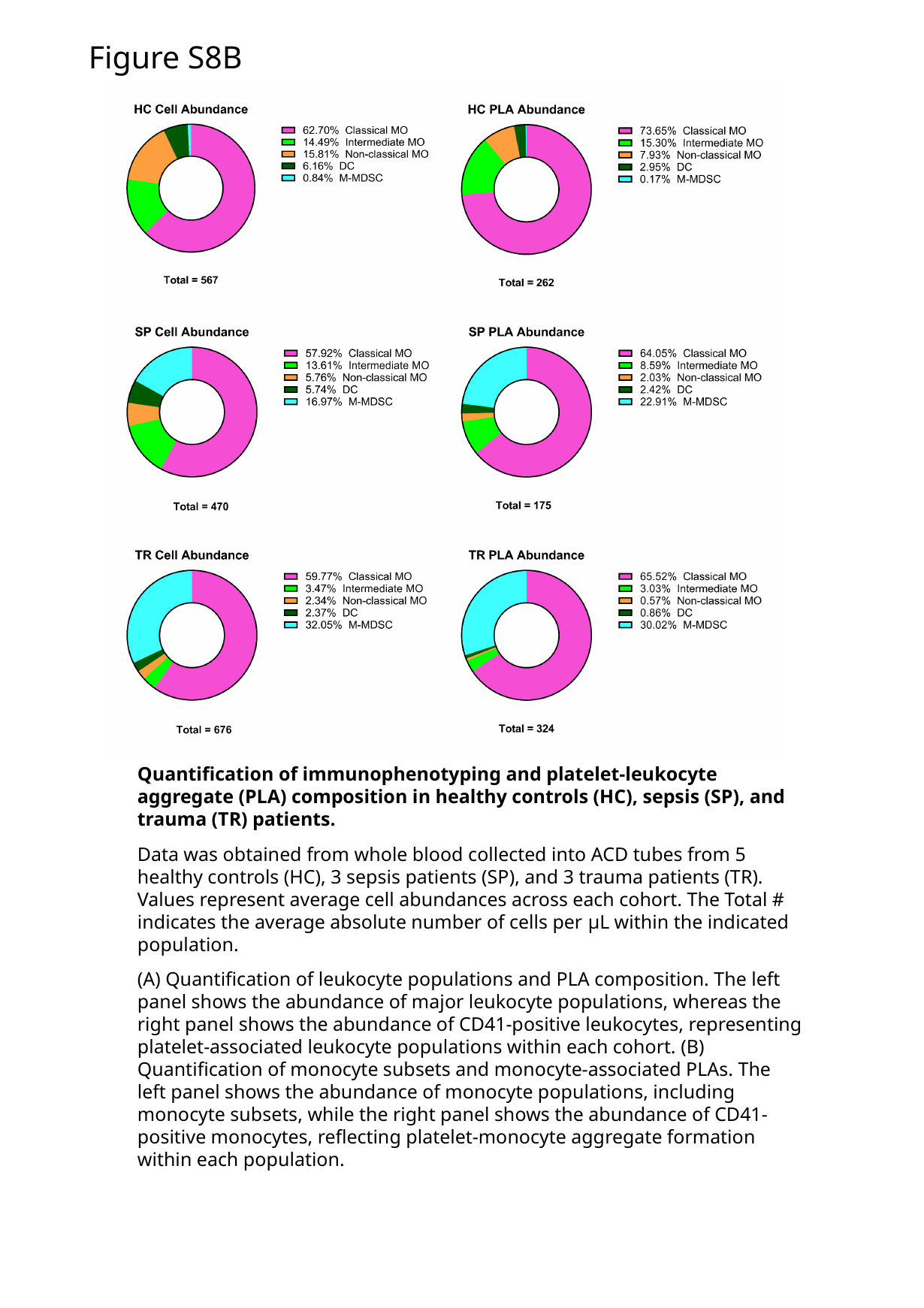

Figure S8B
Quantification of immunophenotyping and platelet-leukocyte aggregate (PLA) composition in healthy controls (HC), sepsis (SP), and trauma (TR) patients.
Data was obtained from whole blood collected into ACD tubes from 5 healthy controls (HC), 3 sepsis patients (SP), and 3 trauma patients (TR). Values represent average cell abundances across each cohort. The Total # indicates the average absolute number of cells per μL within the indicated population.
(A) Quantification of leukocyte populations and PLA composition. The left panel shows the abundance of major leukocyte populations, whereas the right panel shows the abundance of CD41-positive leukocytes, representing platelet-associated leukocyte populations within each cohort. (B) Quantification of monocyte subsets and monocyte-associated PLAs. The left panel shows the abundance of monocyte populations, including monocyte subsets, while the right panel shows the abundance of CD41-positive monocytes, reflecting platelet-monocyte aggregate formation within each population.
